# Physics-Informed Digital Twin Can Predict Cerebral Blood Flow and Cerebral Vascular Regulation Mechanisms in Neurocritical Care Patients

**DOI:** 10.1101/2025.05.22.655581

**Authors:** Jennifer K. Briggs, J.N. Stroh, Soojin Park, Brandon Foreman, Michael M. Tymko, Jay Carr, Melike Sirlanci, Philip N. Ainslie, Richard K.P. Benninger, Tellen D. Bennett, David J. Albers

## Abstract

Cerebral blood flow is vital for brain function and is acutely controlled through a set of physiological mechanisms known as cerebral vascular regulation (CVR)[1–3]. It remains challenging to directly measure the dynamics and function of individual CVR mechanisms[4], limiting our ability to understand and optimize brain perfusion, particularly for neurologically injured patients. Digital twins offer an ideal tool for overcoming this gap because they enable estimation, tracking, and forecasting of unmeasured physiological states[5, 6]. Here, we introduce CereBRLSIM (Cerebral Blood Regulation Latent State Inference and Modeling), a digital twin that integrates physiological knowledge and patient data to infer CVR function and predict cerebral dynamics. Using both *in vivo* experiments and simulated data, CereBRLSIM predicted cerebral hemodynamics with high accuracy and estimated the dynamics of myogenic, endothelial, and metabolic mechanisms underlying CVR. When personalized to neurocritical care patient data, CereBRLSIM differentiated cerebral hemodynamic phenotypes, predicted patient outcomes, and forecasted blood flow with significantly higher accuracy than machine learning models. This work provides a novel, interpretable, and clinically compatible approach for quantifying CVR function and forecasting cerebral blood flow, enabling new opportunities in precision diagnostics and foundational understanding of cerebral hemodynamics.

## 2 Introduction

Cerebral blood flow (CBF) is fundamental for supplying nutrients and clearing waste throughout the brain. Despite comprising a small percentage of total body mass, the brain consumes ≈20% of the body’s oxygen supply. Further, while other organs tolerate 20-40 minutes without blood flow, the brain sustains irreversible damage in under five minutes of inadequate CBF[7, 8]. CBF-related injuries, such as stroke or traumatic brain injury, occur at epidemic levels[9, 10]. Yet, the complex physiological dynamics governing CBF and strategies for its effective management in neurocritical care remain poorly understood. Deeper understanding is hindered by challenges in measuring CBF and the physiological mechanisms governing it in humans.

A promising pathway for overcoming these limitations is the development of mathematical models and digital twins. Mathematical models enable prediction of unmeasured physiological dynamics. Standing alone, mathematical models typically represent average population behavior. Alternatively, these models can be fit to data to represent a unique patient’s physiological function, making the model a “digital twin”. Here, we refer to model fitting as “model personalization”. Digital twins offer the same benefits as stand-alone mathematical models, while also enabling patient-specific predictions. Additionally, when the model is personalized by estimating parameters that have physiological meaning, the digital twin may provide diagnostic capabilities[6, 11, 12]. For these reasons, a digital twin of the cerebral vascular system could be transformative for precision medicine in neurocritical care and enable new insights into cerebrovascular physiology[13, 14].

The cerebral vasculature is highly complex, consisting of a fluid driven by pulsatile pressure gradients through a branching network of compliant tubes whose geometry is heterogeneous across people and actively changes on multiple time scales. Even with continuous measurements of the entire cerebral vasculature, the computational requirements of solving standard fluid dynamics models, such as the Navier-Stokes equations, across the full cerebral vasculature are prohibitive. Alternatively, the problem can be simplified by making assumptions concerning which physical properties dominate fluid flow. The most common simplification is to represent CBF by Poiseulle’s law[15–17]. This can be justified by assuming flow is primarily driven by cerebral perfusion pressure (CPP), created between arterial blood pressure (ABP) and intracranial pressure (ICP), and approximating the vasculature as a single tube with a bulk length and radius (Supp. C.4).

However, predicting CBF with Poiseulle’s law is not straightforward because radius and length are bulk approximations of averaged geometry and cannot be directly measured. Given CBF data, a lumped parameter called “cerebral vascular resistance”[18], could be estimated to represent the effects of vessel geometry. However, if CBF data are not available, vascular resistance cannot be estimated. Using a constant vascular resistance to predict CBF typically fails[19], likely because radius actively changes via cerebral vascular regulation (CVR). Therefore, in the absence of continuous CBF measurements, additional information about radius dynamics is likely essential to model CBF.

CVR is the collective behavior of myogenic, endothelial, and metabolic mechanisms (Fig. 1a)[20–22]. While the term “cerebral autoregulation” is often used to describe the specific response of these mechanisms to changes in pressure[21, 23]. However, here we use the term “CVR” to avoid ambiguity and explicitly refer to all dynamics associated with these mechanisms. Function of CVR mechanisms is heterogeneous across injury and patient demographics[20, 21, 24] and is correlated with neurocritical care patient outcomes[25, 26]. However, the dynamics and function of individual mechanisms are not directly measurable in humans[4].

**Figure 1:**
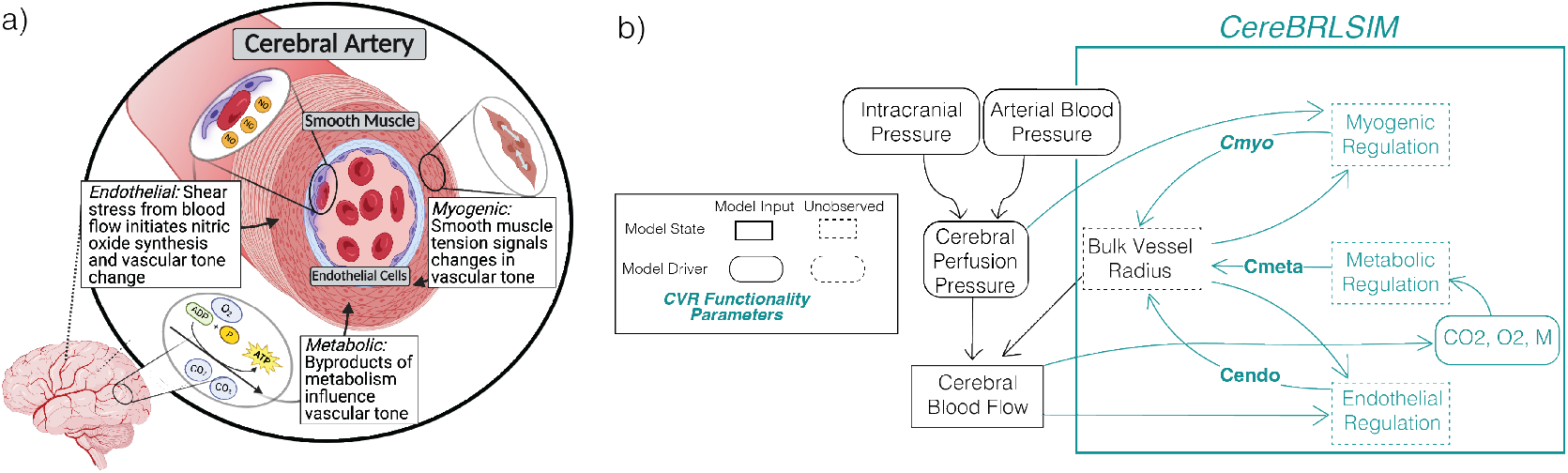
Graphical depiction of CereBRLSIM. **(a)** Cartoon of CVR mechanisms. **(b)** CereBRLSIM includes effects of three CVR mechanisms to change bulk vessel radius. Cmyo, Cmeta, and Cendo mathematically correspond to the functionality of each mechanism.

We hypothesize that a digital twin (called the Cerebral Blood Regulation Latent State Inference and Modeling, or **CereBRLSIM**) that mathematically includes the dynamics of CVR mechanisms with variable functionality on bulk radius in Poiseulle’s equation will enable prediction of heterogeneous CBF and CVR dynamics. To test this hypothesis, we develop the CereBRLSIM and test its predictive ability through three evaluations. First, we show the CereBRLSIM can predict CBF dynamics for a range of in vivo, controlled cerebral hemodynamic experiments. Second, using simulation studies, we show the CereBRLSIM can be personalized by estimating parameters that represent the function of CVR mechanisms, and that these parameters are uniquely estimable. Third, we personalize the CereBRLSIM to neurocritical care data and show that it outperforms other machine learning and deep learning approaches in predicting CBF. Our CereBRLSIM enables novel estimation of CBF dynamics and CVR function, offering new possibilities for cerebral hemodynamics research and the treatment of CBF-related pathologies.

## 3 Results

### 3.1 Cerebral Blood Regulation Latent State Inference and Modeling (CereBRLSIM)

To develop the Cerebral Blood Regulation Latent State Inference and Modeling (CereBRLSIM) model, we approximated global CBF (*Q*) using Poiseulle’s law (eq. 1). This was done approximating the vasculature as a single tube with a bulk length (*L*) and radius (*r*) (Supp. C.4) and assuming CBF if primarily driven by CPP (Δ*P*) and effects of viscosity (*µ*) (Supp. C.3).

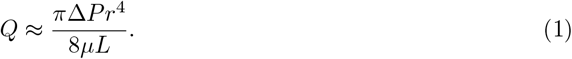

We included effects of myogenic, endothelial, and metabolic mechanisms on bulk radius (*r*) in eq. 1. The myogenic mechanism is triggered by changes in circumferential tension, the endothelial mechanism is triggered by changes in vascular wall shear stress, and the metabolic mechanism is triggered by changes in arteriole blood gases and metabolic activity (Fig. 1a,b). The full model description and equations are given in the methods and mathematical justification is given in Supp. C. Briefly, each CVR mechanism exerts a force (*ϕ*mech) on *r*. This force is modulated by a “CVR parameter” (Cmech), which represents the ‘functionality’ of each mechanism (Cmech). These CVR parameters will later be estimated with patient data to predict mechanism functionality. The work (change in radius per unit force) done by that mechanism is modulated by a length tension function (*K*(*r*)), which represents the length tension relationship of smooth muscles[27, 28].

### 3.2 CereBRLSIM Reproduces Data from *in vivo* Experiments

We verified CereBRLSIM using data from *in vivo* experiments. We first used three studies that assess vascular response to dynamic changes in physiology (occurring within minutes). These experiments were: (Fig. 2a) a dynamic cerebral autoregulatory challenge, typically associated with myogenic regulation[29, 30]; (Fig. 2b) a flow-mediated dilation test, associated with endothelial-dependent regulation[31, 32]; and (Fig. 2c) a neurovascular coupling test, associated with metabolic regulation[33].

**Figure 2:**
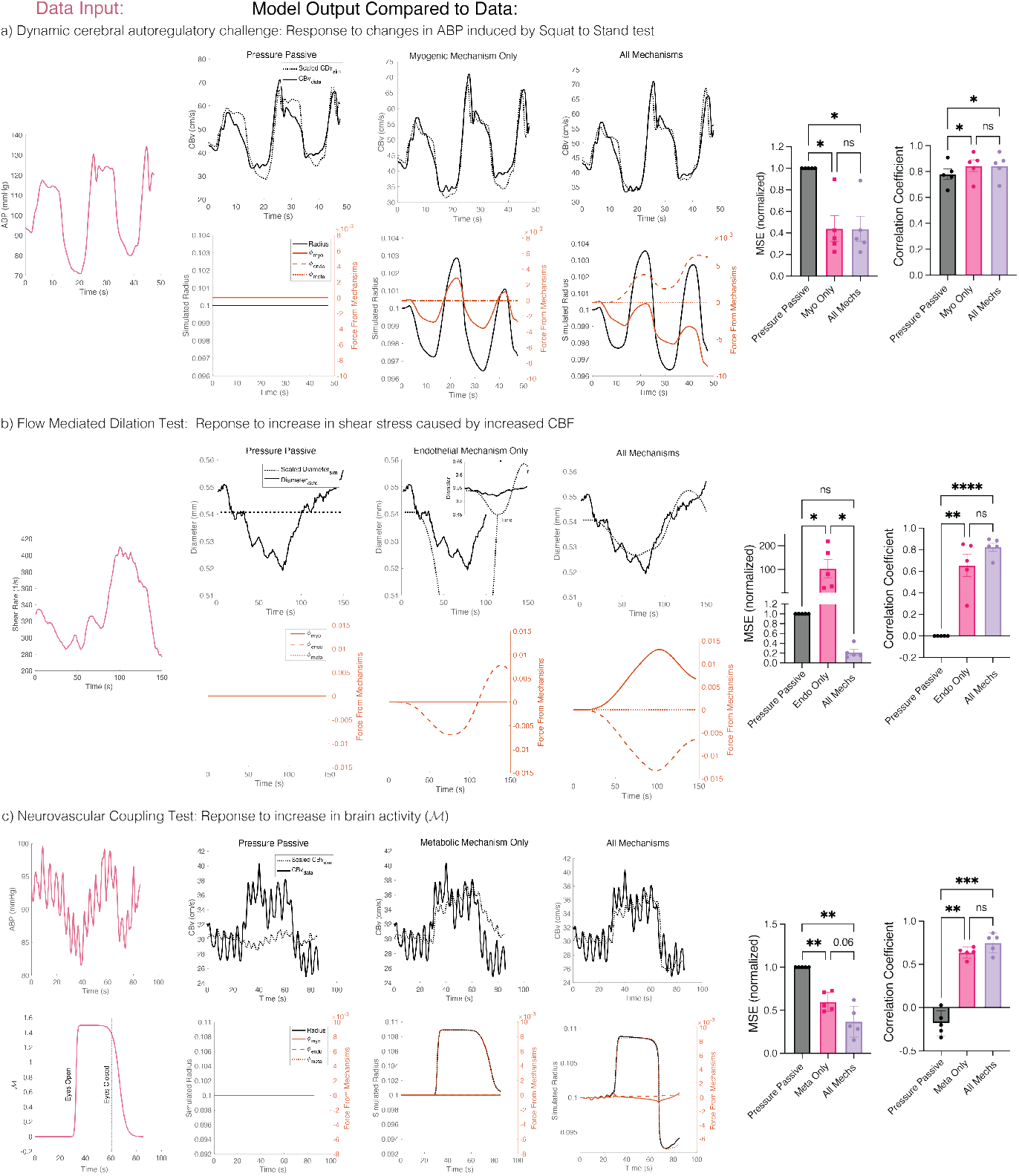
CereBRLSIM Closely Predicts *In Vivo* Experiments. Three *in vivo* experiments with five patients each were used for model validation: **(a)** Squat to stand test, primarily associated with myogenic mechanism, **(b)** Flow mediated dilation test, primary associated with endothelial mechanism, **(c)** Neurovascular coupling test, primarily associated with metabolic mechanism. Pink time course shows data used to drive the model. Patient data (solid black line) were compared the pressure-passive simulation (no CVR) (dotted line, left), simulation of only the primary CVR mechanism (dotted line, center), and all three CVR mechanisms (dotted line, right). Lower plots show CereBRLSIM-predicted radius (black line) and the virtual force from each mechanism (*ϕ*_*mech*_, orange lines). Bar plots show quantification of mean square error (MSE) normalized to the pressure passive simulation and correlation coefficient between model and patient data. Dots indicate model performance for each patient. All statistical tests are one way ANOVA with Sidak multiple comparison. *P<0.5, **P<0.01, ***P<0.001, ****P<0.0001.

During a dynamic cerebral autoregulatory challenge, ABP was perturbed by instructing subjects to transition from squatting to standing every ten seconds (Fig. 2a, pink). In the flow-mediated dilation tests shear stress was increased by inhaling CO_2_ to dilate upstream vessels (Fig. 2b, pink). Shear stress was calculated as 4 x Peak envelope velocity / vessel diameter measured using duplex ultrasound. During the neurovascular coupling test, cerebral activity was increased through an eye-opening and reading task (Fig. 2c, pink).

To verify CereBRLSIM’s ability to reproduce vascular dynamics from these *in vivo* experiments, we simulated eq. 1 with unchanging radius by setting CVR parameters: Cmyo, Cendo, and Cmeta to zero (Fig. 2, “Pressure-Passive”). We term this the term “pressure-passive”, drawing from clinical terminology used to describe a patient presumed to have impaired vascular reactivity[34–37]. Importantly, the term has broader clinical applications beyond our usage here. Second, to test if vascular dynamics could be predicted by simulated only the primary mechanism associated with each *in vivo* test, we turned “off” the non-primary mechanisms by setting their CVR parameters to 0 and turned “on” the primary mechanism by setting its CVR parameter to 1 (Fig. 2, “mechanism” only). For example, in the dynamic cerebral autoregulatory challenge, the myogenic mechanism is the primary mechanism. Therefore, Cmyo=1 while Cendo=Cmeta=0. Finally, to test if CereBRLSIM could predict vascular dynamics when all mechanisms were functional, we turned all mechanisms “on” by setting Cmyo=Cendo=Cmeta=1 (Fig. 2, All Mechanisms).

Predictive ability was quantified using mean square error (MSE) and correlation coefficient (Corr) between the data and model output. Raw MSE is reported in the Table. A.1 and Fig. A.4, while normalized MSE is reported in Fig. 2. Because CereBRLSIM emulates the global cerebral vasculature, whereas experimental measurements correspond to individual arteries, CereBRLSIM simulation (Fig. 2 black, dotted line) was scaled before comparison. Scaling was determined through linear regression between the full CereBRLSIM and patient data and was held constant across all experimental conditions. However, our findings were consistent when scaling was calculated separately for each model condition (Fig. A.4).

For the dynamic cerebral autoregulatory challenge, simulated cerebral blood velocity (CBv_sim_), was compared to CBv measured by transcranial doppler. Simulations using only the myogenic mechanism and all mechanisms had lower MSE and higher correlation compared to the pressure passive model (Fig. 2a and Fig. A.1). In the flow-mediated dilation test, simulated diameter was compared to diameter measured by transcranial doppler. By definition, pressure passive simulations did not predict a change in vessel diameter (Fig. 2b, left). When simulating only the endothelial mechanism, the predicted change in diameter was much larger than experimentally measured (Fig. 2b, Endothelial Mechanism Only). Simulations using all mechanisms had lower MSE and higher correlation than both the pressure-passive and the endothelial-only model (Fig. 2b, right and Fig. A.2). For the neurovascular coupling test, brain metabolic state (*ℳ*) was simulated as a step function that increased directly after eyes were opened and decreased slowly when the eyes were shut (Fig. 2c, pink). Simulated cerebral blood velocity was compared to CBv measured by transcranial doppler. Again, the pressure-passive simulations were least reflective of patient data. Modeling only the metabolic mechanism predicted the observed increase in CBv but showed a delayed response to the decrease in CBv compared to patient data. Simulations using all mechanisms were the most accurate in predicting experimental data (Fig. 2c, right and Fig.A.3). These results indicate that CereBRLSIM was strongly predictive of all experiments and that all CVR mechanisms are helpful in explaining vascular dynamics.

We also assessed CereBRLSIM’s ability to *in vivo* predict experiments involving slower (>10 min) and larger changes in end tidal CO_2_ (ETCO_2_) and CPP[38, 39], both of which show a non-linear relationship with CBF. We hypothesized that this non-linear relationship could be explained by the length-tension relationship (*K*(*r*)), which describes the physical limitations of vessel size and contractility. When the length-tension relationship was not mathematically accounted for, CereBRLSIM did not reproduce the known relationships between CBF and ETCO_2_ (Fig. 3a) or CPP (Fig. 3d). However, when the length-tension relationship was accounted for by simulating *K*(*r*) with eq. 9 (Fig. 3b,e), CereBRLSIM closely predicted experimental data[38, 39] (Fig. 3c,f). Importantly, these results emerge without introducing additional mathematical nonlinearities within individual CVR mechanisms. These findings support the hypothesis that the non-linear relationships between CBF, ETCO_2_, and CPP can be explained by the length-tension relationship of vessels.

**Figure 3:**
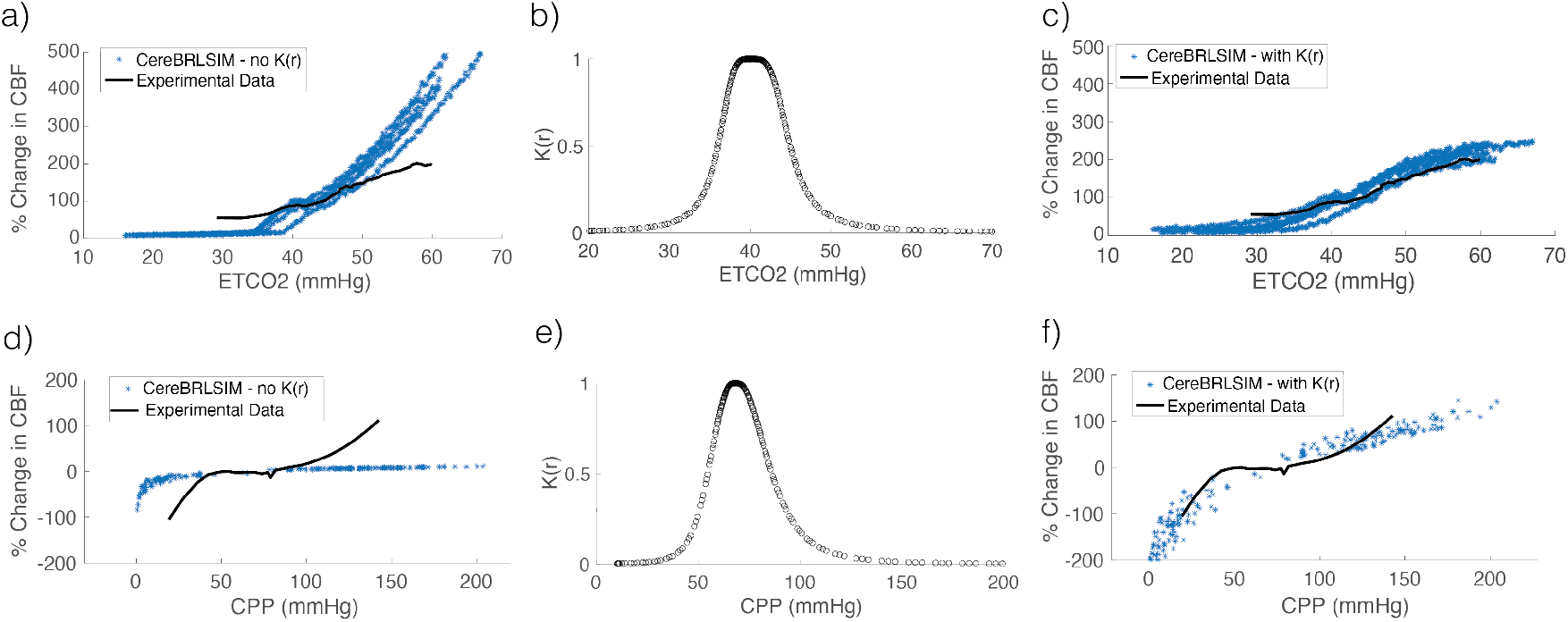
Length Tension Function Can Predict Nonlinear CBF Relationships. **(a)** Change in CBF (blue) *without* length tension function (*K*(*r*)) compared to experimental data (black). **(b)** Relationship between ETCO2 and *K*(*r*). **(c)** Simulated change in CBF *with* (*K*(*r*)). **(d-f)** as in a-c but comparing relationship between CBF and CPP.

Together, these results verify that CereBRLSIM quantitatively and qualitatively recapitulates experiments of cerebral vascular tone regulation.

### 3.3 CVR Parameters are Uniquely Estimable

CereBRLSIM contains three CVR parameters: Cmyo, Cendo, and Cmeta, which correspond to myogenic, endothelial, and metabolic function, respectively. For example, after an increase in CPP and a corresponding rise in CBF, a functional myogenic response (Cmyo=10) will lead to a rapid decrease in CBF, while an absent myogenic response (Cmyo=0) shows no restoration of CBF (Fig. 4a). CereBRLSIM was personalized by estimating these parameters with patient data. We conducted simulation experiments to assess estimation certainty and accuracy. First, we simulated nine phenotypes with variable CVR parameters using CPP and ETCO_2_ data from a traumatic brain injury patient (Fig. 4b). The phenotypes included different combinations of CVR parameters being ‘on’ or ‘off’ (1 or 0) or being partially ‘on’ (0.5, 0.5, 0.5).

**Figure 4:**
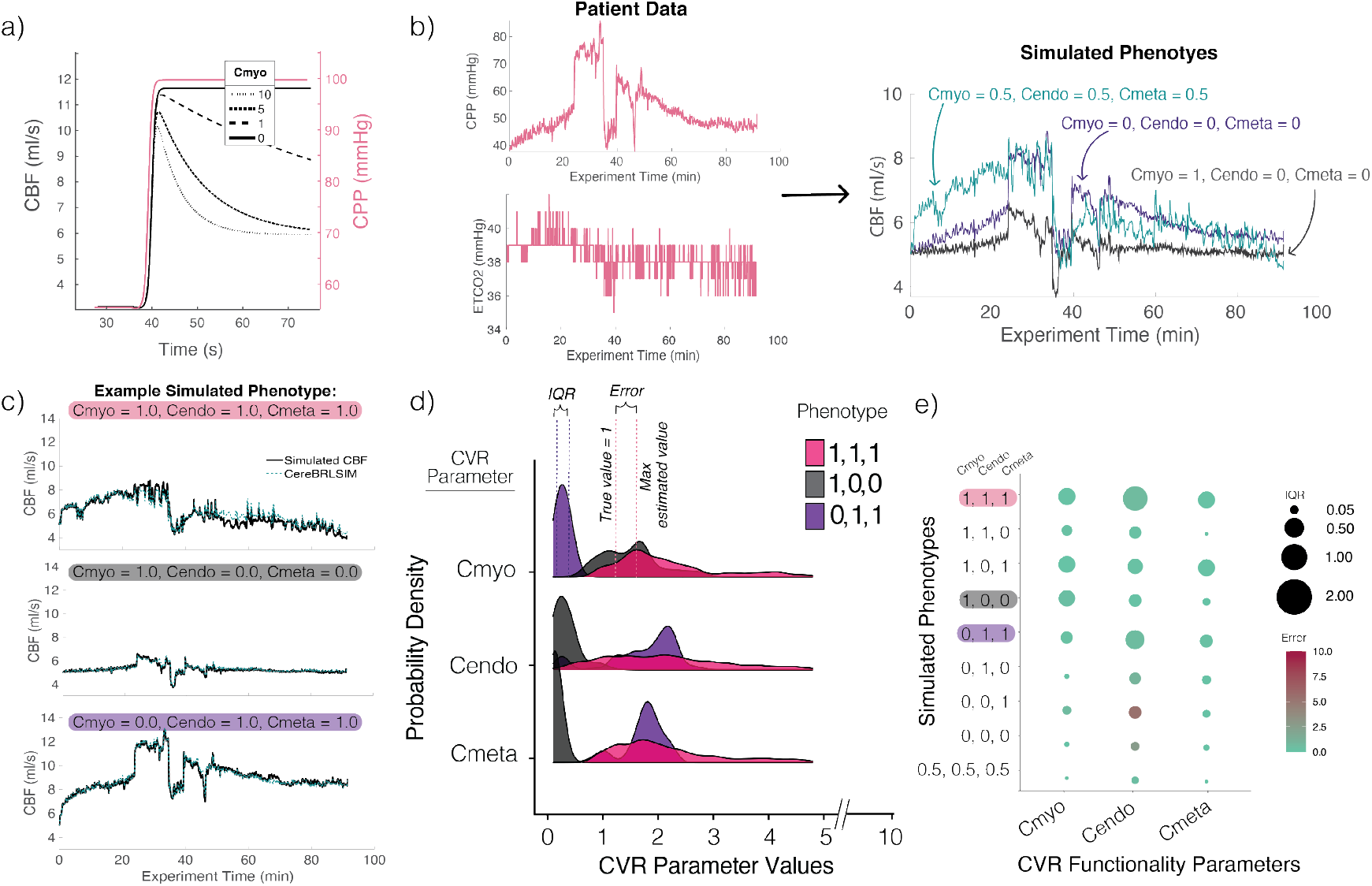
CVR Functionality Parameters Are Uniquely Estimable. **(a)** Example showing how Cmyo impacts CBF response (black) to an increase in CPP (pink). **(b)** Patient data (pink) used to simulate CBF phenotypes (three representative phenotypes are shown on the right). **(c)** Example of true CBF and predicted CBF by CereBRLSIM (teal) for three example phenotypes. **(d)** Probability distributions of CVR parameters for example patients corresponding to c. **(e)** Quantification of interquartile range (dot radius) and error (dot color) of estimated CVR functionality parameters for nine simulated phenotypes.

We estimated CVR parameters with uncertainty using Markov Chain Monte Carlo (MCMC) for each of the simulated CBF phenotypes. Figure 4c shows three example phenotypes (black) and CereBRLSIM-predicted CBF (teal). For all cases, CereBRLSIM closely predicted CBF. We quantified uncertainty and error for each parameter estimate. Figure 4d shows parameter distributions for patients corresponding to Figure 4d. Generally, Cmyo, Cendo, and Cmeta were estimated with relatively low error and high certainty (Fig 4e). The MSE (normalized to the estimation range 0-10) was: MSE Cmyo=1.57±1.77%, Cendo=5.28±17.3%, Cmeta=0.88±0.07%. The average interquartile range (IQR) of the estimated posterior distribution was: Cmyo=0.61±10.5%, Cendo=4.12±13.87%, Cmeta=0.972±10.3%. When Cmyo and Cendo were absent and Cmeta was functioning, Cendo was poorly estimated, indicating that Cendo may only be reliable when Cmyo is at least slightly functional (e.g. not equal to 0).

The results were similar when variables used to drive the model were held fixed (Sup. Fig A.5a,b) or a single parameter was estimated while the other two were held fixed at their true values (Sup Fig A.5c). These results indicate that when data are sufficiently descriptive (e.g. change over time), CVR parameters can be estimated with generally high accuracy and certainty.

### 3.4 CereBRLSIM Predicts CBF and CVR function for Traumatic Brain Injury Patients

Next, we validated CereBRLSIM in neurocritical care data. Because the magnitude of CBF is dependent on the location of the perfusion monitor, we estimated a scaling parameter (*α*) (eq. 8, Supp. C.6, Table A.2) for the first ten minutes of patient data. We then used MCMC to estimate CVR parameters over an hour of patient data (Fig. 5a). We assessed if: (1) the personalized CereBRLSIM can predict CBF and (2) CVR parameters can differentiate pressure-flow phenotypes and patient outcomes. These phenotypes, as defined by[19], are: positive, indicated by a positive global correlation between CBF and CPP; zero, indicated by a near-zero correlation; and negative, indicated by a negative correlation (Fig. 5b). Patients with functional CVR are assumed to show no correlation between CBF and CPP.

**Figure 5:**
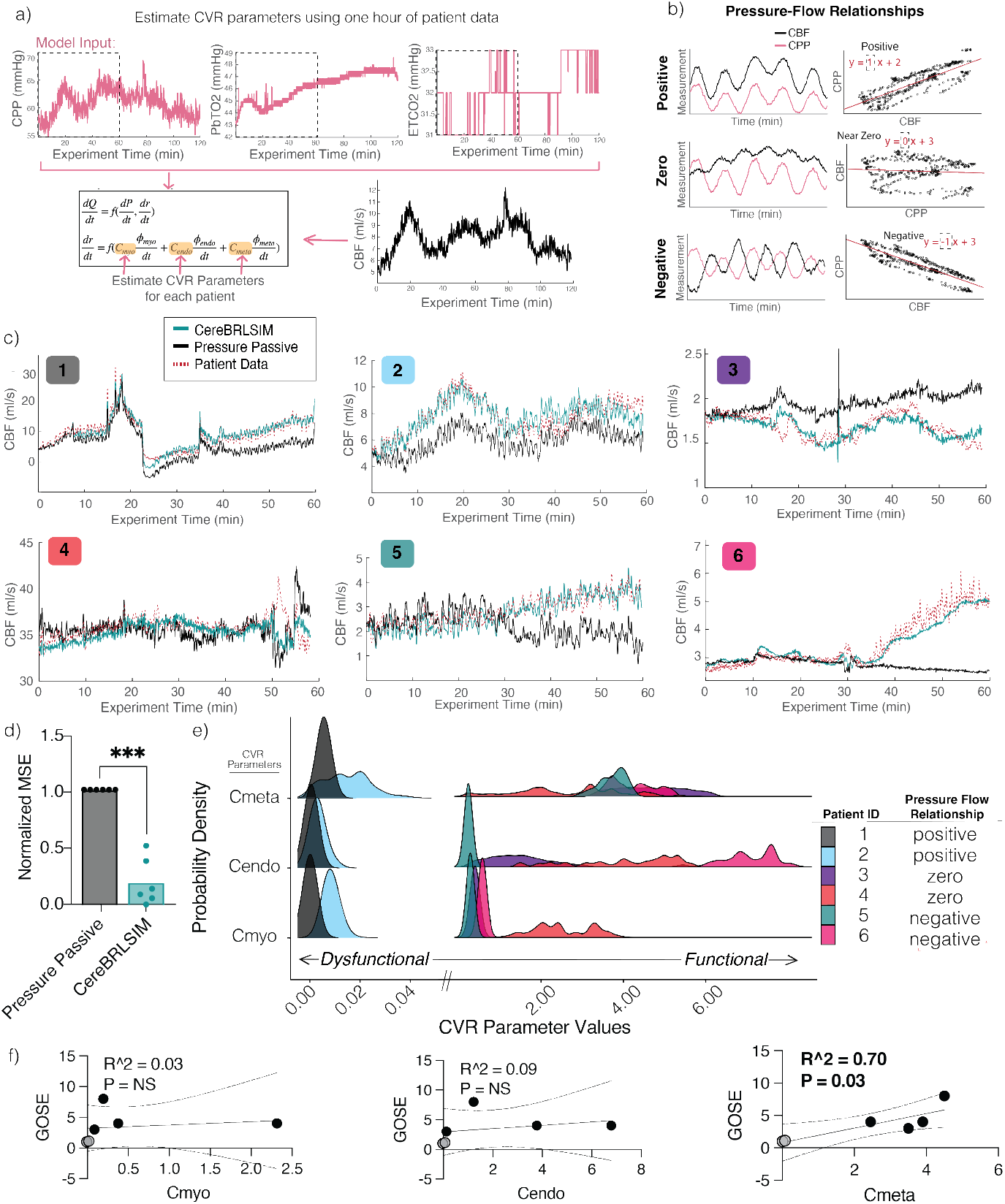
CereBRLSIM Predicts Cerebral Blood Flow and Estimates CVR Mechanism Functionality. **(a)** Methodological outline for personalized CereBRLSIM. MCMC identifies CVR functionality parameter posterior distributions given one hour of patient data. **(b)** Example of three pressure flow relationship (PFR) phenotypes. Left plots show hypothetical CBF and CPP time series. Right plot shows linear correlation between CPP and CBF[19]. **(c)** Qualitative comparison between CereBRLSIM (teal), pressure-passive simulation (black) and patient data (red). **(d)** Quantification mean square error (MSE) normalized to the pressure-passive assumption. P value is given by a paired students t-test P*<*0.001. **(e)** Probability density functions of each estimated CVR functionality parameter for each patient. Colors and Patient ID’s correspond to (c). Lower parameter values indicate less functional CVR. **(f)** Correlation between average CVR functionality parameter estimates and Glasgow Outcome Scale-Extended (GOSE). Grey dots indicate patients with 0 GOSE and positive pressure flow relationships.

Visual inspection shows high concordance between CereBRLSIM-predicted CBF and patient data (Fig. 5c). CereBRLSIM also had lower MSE than the pressure-passive model for all patients (Fig. 5d, CereBRLSIM MSE: 0.31±0.59 vs Pressure Passive MSE: 2.72±4.04), indicating that personalized CVR is necessary to predict CBF in neurocritical care patients. Further, CVR parameters distinguished positive pressure-flow phenotypes from negative or zero pressure-flow phenotypes (Fig. 5e). Patients with positive pressure flow relationships had much lower CVR parameter values (corresponding to dysfunctional CVR) than those with negative or zero. To assess if the CVR parameter estimates were informative of known outcome metrics, we used the Glasgow Outcome Scale-Extended (GOSE), which quantifies patient neurological state six months after discharge. GOSE ranges from 0-8 with 0 indicating deceased and 8 indicating complete neurological recovery. Cmyo and Cendo were not linearly related to GOSE, but Cmeta was linearly correlated with GOSE (Fig. 5f). Therefore, Cmeta was able to predict patient outcomes, particularly very poor outcomes (GOSE=0) (with the caveat of small sample size). Similar results were found when correlating CVR vascular metrics with the Pressure-Reactivity Index (PRx) and Mean Flow Index (Mx), which are time-series based metrics of cerebral autoregulation (Supp. A.6). In our data set, PRx and Mx were not predictive of GOSE. Therefore, CereBRLSIM can be personalized to neurocritical care data to predict CVR function.

### 3.5 CereBRLSIM Forecasts CBF with High Accuracy

Finally, we assessed the forecasting ability of CereBRLSIM. For each patient, we personalized CereBRLSIM using their estimated CVR parameters and then simulated CBF forward one hour (Fig. 6a).CereBRLSIM took 1.1±0.42 seconds to forecast one hour of patient data, indicating it can be solved much faster than real time. By construction, CereBRLSIM hypothesizes the change in the bulk radius (Fig. 6b), and the action of each CVR mechanism over time (Fig. 6c). To assess how CereBRLSIM performed compared to traditional machine learning methods, we also trained a neuralODE (a continuously differential neural network that can be simulated forward in time), and a multivariate linear regression model with the same one hour of data given to CereBRLSIM and then forecasted these models forward in time. Compared to the pressure-passive assumption, neural ODE, and multivariate regression, CereBRLSIM predicted CBF with the lowest MSE at 5, 10, 30, and 60 minutes into the future (Fig. 6d, Table. A.3). These results indicate that CereBRLSIM can be personalized to neurocritical care patient data to predict CVR function and CBF dynamics with strong fidelity.

**Figure 6:**
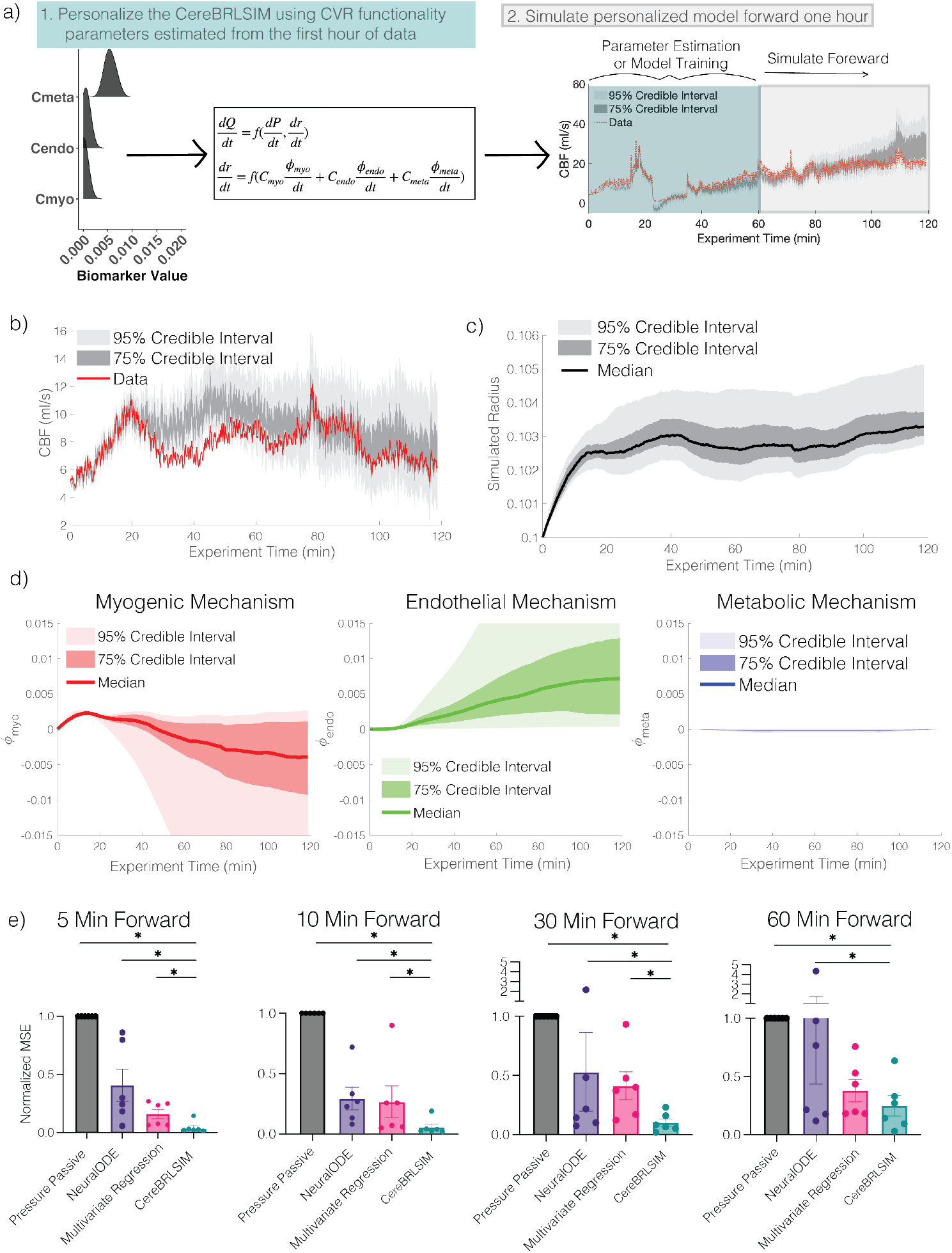
CereBRLSIM Predicts CBF with Higher Accuracy than Other Machine Learning Models. **(a)** Methodological outline: After CereBRLSIM was personalized to one hour of data, it was simulated forward an additional hour. **(b)** Representative forecast. **(c)** Representative bulk radius. **(d)** Representative virtual forces from each CVR mechanism. **(e)** Quantification of CBF forecasts from three models normalized to the pressure-passive model.

## 4 Discussion

Adequate CBF is essential for brain function and is tightly regulated by CVR mechanisms. CBF-related injuries, like Stroke and traumatic brain injury, impact CVR and CBF, thereby impacting patient outcomes. However, measurements of CVR function and dynamics are not feasible in clinical settings and measurement of CBF is rare[40]. A digital twin that can quantify CVR function and forecast CBF could be transformative for the study and treatment of CBF-related injuries by allowing for personalized diagnostics, virtual experiments to identify optimal treatments, and real-time monitoring of physiological function that is otherwise difficult to measure. In this study, we developed a novel digital twin of the cerebral vascular system, Cerebral Blood Regulation Latent State Inference and Modeling (CereBRLSIM), that can be estimated with neurocritical care patient data, opening new opportunities in CBF-related research and clinical decision support.

We conducted three evaluation steps to test the performance of CereBRLSIM. First, we showed CereBRLSIM closely reproduced dynamics of five in vivo cerebral hemodynamic experiments (Figs. 2 and 3). Next, using simulated data, we showed that CVR parameters, which correspond to mechanism function, could be uniquely estimated (Fig. 4). Finally, we personalized CereBRLSIM to six neurocritical care patients and showed that it was able to forecast heterogeneous CBF dynamics with lower MSE than machine learning or deep learning models and that CVR parameters could differentiate patients based on their subsequent outcomes and pressure flow phenotypes (Figs. 5 and 6).

The definition and grouping of mechanisms underlying CVR is variable[20, 21, 41]. Here, we group mechanisms based on time scales and physiological triggers to mathematically resolve their dynamics. Terms “cerebral vascular regulation”, “cerebral vascular reactivity”, and “cerebral autoregulation” are also used to describe these mechanisms or their subsets. To avoid ambiguity, we use the term cerebral vascular tone regulation to refer to the collective effects of all three mechanisms.

Our results show that modeling individual mechanisms is not usually sufficient to predict dynamics of in vivo experiments (Fig. 2). This aligns with prior studies indicating such experiments may not isolate mechanism dynamics[4, 21, 22, 42]. CereBRLSIM may help address this gap because it can estimate individual CVR mechanism function (as in Fig 5) and model separable dynamics of each CVR mechanism (Fig 2, Fig 6c,d).

It is well-established that CBF demonstrates non-linear relationships with CO_2_[38, 43] and CPP[44]. Our results indicate these non-linear relationships can be explained through the length-tension relationship of smooth muscles and do not require non-linearities or ranges of regulation to be directly encoded in the CVR mechanisms themselves. Future research could test this hypothesis experimentally and assess whether asymmetries in the *K*(*r*) curve could explain asymmetric responses to CPP[39, 45].

Currently, there is no established method to estimate CVR function mechanistically in neurocritical care patients. This makes CereBRLSIM potentially impactful, but also poses challenges validating if estimated CVR parameters correspond to their intended physiological meaning. While we showed that CVR parameters are uniquely estimable, which is important from a mathematical and optimization perspective (Fig. 4), that does not confirm their physiological interpretation. Proper validation of CVR parameters would involve human or animal experiments in which each CVR mechanism is blocked or genetically knocked-out[46]. This remains a topic of future work. However, three of our findings support the predictive value of CVR parameters. First, by estimating CVR parameters, CereBRLSIM could reproduce CBF dynamics (Fig. 5c) better than the pressure passive model. This suggests that our mathematical representations of CVR reasonably capture CBF dynamics and supports the claim that CVR parameters may reflect underlying mechanism function. Second, CVR parameters were predictive of impaired CVR, as suggested by positive pressure-flow relationships (Fig. 5e). Third, CVR parameters correlate with patient outcome (Fig. 5f). Together, these findings support the interpretation of CVR parameters and suggest their potential use as computational biomarkers for clinical decision support.

Digital twins fit into a broader field of physics-informed machine learning, where models are informed by data but constrained by fundamental principles of physics and physiology. CereBRLSIM highlights the potential benefits of physics-informed machine learning because it offers interpretability and also improved prediction accuracy compared to a more traditional deep learning approach (e.g., neural ODE, Fig. 6).

### 4.1 Limitations and Future Work

CVR mechanisms are thought to operate on different spatial scales[20]. Representing the vasculature using a single bulk vessel (Supp. C.4), limits our ability to resolve these scales. This choice was necessary to resolve CVR parameters given data sparsity. Future work could incorporate spatially resolved data (e.g., MRI) into a spatially extended version of CereBRLSIM to examine regional CVR and CBF dynamics.

CVR function likely changes over time, especially in neurocritical care settings. In our experiments, we assume stationary CVR parameters to estimate and forecast CereBRLSIM. This assumption appears justified because of the strong predictive performance. However, future work could use filtering methods to update CVR parameters dynamically, enabling real-time tracking of patient state, which may offer valuable clinical insight.

In non-pathologic conditions, smooth muscles are typically able to maintain vessel radius on timescales longer than tens of seconds, making the impact of vascular compliance negligible on longer timescales[16]. Therefore, in this study, we addressed the impact of vascular compliance on short time scales by applying a smoothing average (Supp. C.5 and C.6). However, if isometric contraction is impaired, which may occur with CVR impairment, the radius will passively change with pressure over all time scales. Although CereBRLSIM does not explicitly model this passive relationship, we account for its effects by estimating *α*, the parameter controlling pressure-flow scaling (Supp. C.6). Patients with impaired isometric contraction should have larger *α*values. Indeed, patients with positive pressure-flow relationships, indicating no CVR and potentially impaired isometric contraction, exhibited higher *α*values than those with negative or zero pressure-flow relationships (Table A.2). Thus, while the simulated radius loses physiological interpretability when isometric contraction is impaired, the model remains useful for predicting CBF because *α*implicitly, albeit crudely, captures the impact of vessel compliance on CBF. Alternative approaches to account for vessel compliance include using a Windkessel model[47–49], which partitions vessel geometry into “compliant” and “rigid” components (Supp. C.7). This partitioning is not suitable in our case because geometry is a core consideration of the model. Future work should incorporate heterogeneous effects of compliance on *r*.

CVR mechanisms rely on physiological triggers that are not directly measured (e.g., brain metabolic activity). CereBRLSIM uses available multimodal data to approximate these triggers (see methods). Future work should explore whether these triggers can be approximated using alternative or improved data sources, or estimated through machine learning approaches.

We estimated CVR parameters with uncertainty quantification using Markov Chain Monte Carlo estimation (MCMC). Given that the model was run in finite time, the estimated posterior distributions of the CVR computational biomarkers are not exact. To minimize this problem, we simulated multiple chains and methodically chose chains that converged to similar posterior distributions. Still, for some patients, some simulations run from the same posterior displayed very different dynamics than the optimal solution. We do not believe this impacts the findings except that the uncertainty in posterior distributions may be over-estimated. Future work should explore other MCMC methods to improve convergence and reduce uncertainty in posterior distributions.

## 5 Conclusion

In this study, we developed and validated a novel digital twin of cerebral hemodynamics that includes three mechanisms underlying cerebral vascular regulation (CVR). The Cerebral Blood Regulation Latent State Inference and Modeling digital twin (CereBRLSIM) can be personalized to patient data by estimating the parameters corresponding to the CVR mechanism function. We showed that CereBRLSIM was able to predict data from controlled experimental with high accuracy, estimate the function of CVR mechanisms in simulation studies, and predict CVR mechanisms and forecast cerebral blood flow in real-world neurocritical care patient data. By enabling real time monitoring of patient state, tracking of currently difficult-to-measure CVR mechanism dynamics, and forecasting of CBF, CereBRLSIM enables new avenues for scientific investigations and precision medicine in cerebrovascular health.

## 7 Methods

### 7.1 CereBRLSIM: A Digital Twin of Cerebral Vascular Hemodynamics and Regulation

To construct a model of bulk cerebral hemodynamics that includes CVR mechanisms (Fig. 1b), we assume that global CBF can be approximated as Poiseulle flow through a single tube (eq. 1) with bulk radius (*r*) and length (*L*). Accounting for the effects of vascular compliance by passing a smoothing average over pressure (*P*), flow (*Q*), and *r* and assuming changes in viscosity (*µ*) and *L* are negligible (Supp. C.5), the derivative of eq. 1 with respect to time is:

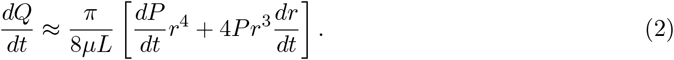

The total work (or change in radius/unit of force) done by vascular smooth muscles is dependent on the radius. This is known as the length-tension, or radius-tension, relationship[1, 2]. We represent this relationship through a length-tension function (*K*(*r*)), which governs the extent to which CVR mechanisms can change vessel radius. Vessel radius is actively controlled by myogenic, endothelial, and metabolic mechanisms. We define a ‘virtual force’ by each mechanism 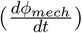,which describes the mechanism’s response to external triggers. This force is ‘virtual’ because it abstracts away the physical force exerted by smooth muscles and does not account for work against transmural pressure. The total change in radius is the sum of CVR virtual forces multiplied by *K*(*r*).

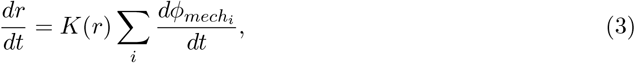

where *ϕ*_*mech*_ = *{ϕ*_*myo*_, *ϕ*_*endo*_, *ϕ*_*meta*_*}* (Fig. 1b, teal).

#### CVR Mechanisms

Virtual forces from each mechanism 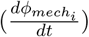 are modeled using a consistent mathematical expression with four components:

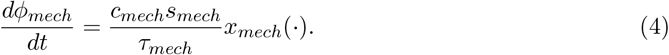

The first component, *c*_*mech*_, is a scaling factor relating to the *function* of each mechanism, which will later be estimated to personalize the digital twin to patient data. We call this quality the ‘CVR parameter’. The second component, *s*_*mech*_, is a normalization factor that ensures the mechanisms are comparable in scale and maintains the correct units. The third component, *τ*_*mech*_, describes the distinct time scales of each mechanism. Values for *τ*_*mech*_ were determined via the literature[3] and manually adjusted to match experimental data (Fig. 2 and Supp. B.1). The final component, *x*_*mech*_(*·*), is the response of each CVR mechanism to physiological triggers.

#### Myogenic Mechanism Response

The myogenic mechanism involves the vascular response to changes in arteriole wall tension, typically due to changes in transmural pressure. Changes in tension triggers stretch-sensitive ion channels that initiate vasomotion[4, 5]. Vascular wall tension is a function of the vessel radius and pressure (*f*_1_(*r, P*)) (defined in Methods: Model Implementation eq. 10). Therefore, the myogenic response is modeled as the deviation of wall tension (*f*_1_(*r*(*t*), *P* (*t*))) from basal tension (*f*_1_(*r*_*opt*_, *P*_*opt*_)):

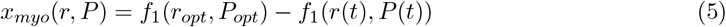

#### Endothelial Mechanism Response

The endothelial mechanism describes when shear stress triggers the endothelium to produce vasodilators (primarily nitric oxide (NO)[6]). Shear stress is given by: 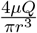 [7]. After combining constants 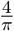 into *S*_*endo*_, the deviation of shear stress from baseline is: 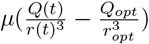.Experimental studies show that shear-mediated dilation is temporally delayed by *tau*_*endo*_ after an increase in shear stress[6, 8]. Therefore, a delayed differential equation is required to represent the endothelial-derived force:

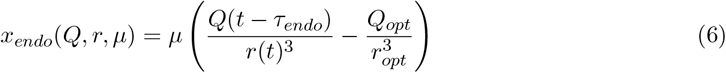

#### Metabolic Mechanism Response

The metabolic mechanism ensures that blood flow is adequate to meet metabolic demand and clear waste. Physiological processes underlying the metabolic mechanism are abundant and incompletely understood. Neurovascular coupling (NVC) and reactivity to blood gases, including partial pressure of carbon dioxide (pCO_2_) and partial pressure of oxygen (pO_2_), and are thought to be the primary metabolic related-contributors to CVR[4]. NVC is often considered an independent mechanism. However, because NVC acts on a similar timescale and has similar emergent dynamics as CO_2_ reactivity (e.g., increases in metabolism cause vasodilation), it is difficult to distinguish the effects without direct experimentation and is therefore modeled in the same mechanism to avoid model estimability issues. Therefore, the metabolic mechanism is modeled as the sum of three components: vasoreactivity to partial pressure of carbon dioxide (*pCO*_2_), vasoreactivity to partial pressure of oxygen (*pO*_2_), and neurovascular coupling (NVC)[4].

#### Vasoreactivity to Carbon Dioxide

The emergent relationships between CBF and carbon dioxide are nonlinear [9]. We hypothesize (Fig. 3) that this nonlinearity can be explained by the length-tension function (*K*(*r*)) rather than a non-linear metabolic response to CO_2_ levels. Therefore, we assume 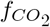 is linearly related to partial pressure of carbon dioxide [4].

#### Vasoreactivity to Oxygen

The cerebral vasculature is only reactive to oxygen cases of severe hypoxemia[4, 10–12]. Therefore, we 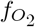 as sigmoid dependent on PO_2_ with some ischemic threshold *γ*.

#### Vasoreactivity in relationship with neurovascular coupling

Assuming resting radius is accompanied by resting metabolic state, NVC is modeled as the change in cerebral metabolism 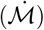.

The metabolic mechanism response is modeled as the sum of these three components:

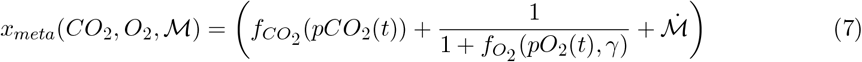

Together, eqs. 2 - 7 define the CVR model:

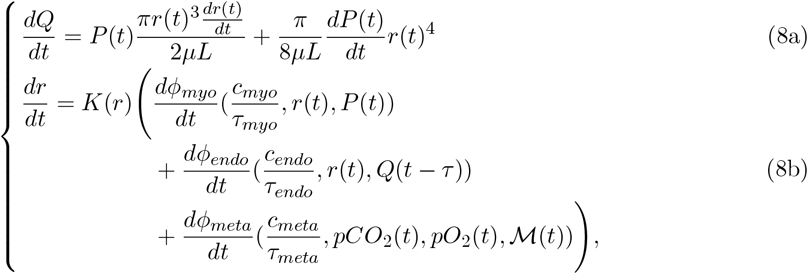

where pressure *P, pCO*_2_, *pO*_2_, and 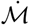 are driven by external data inputs. All parameter values are in Table B.1 and the full set of model equations as they are solved with patient data are in the Supp. B.

### 7.2 Model Implementation

All simulations were conducted in MATLAB, and the CVTR model was solved using Runge-Kutta (2,3) with fixed time steps.

#### Length-tension function

The length-tension function (*K*(*r*)) was modeled as a Gaussian centered at *r*_*opt*_. A small constant (10^−8^) is added to *K*(*r*) to ensure the radius does not get stuck in an over dilated or over constricted position:

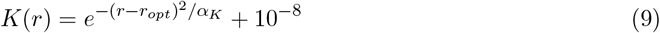

#### Myogenic tension function

*f*_1_ was empirically chosen to best predict experimental evidence, which suggests that smooth muscle response increases exponentially with deviations in resting radius (*r*_*opt*_) and linearly with deviations from resting pressure (*P*_*opt*_)[13, 14]. This behavior is approximated by eq. 10, which was the lowest order polynomial that predicted.

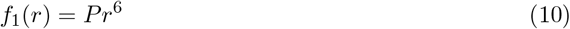

#### Endothelial Mechanism Delay

In eq. 8, the endothelial mechanism is written as a delayed differential equation. To avoid the computational limitations imposed by a delayed differential equation, we model the endothelial delay response using the linear chain trick, a common mathematical technique in which sub-states (*h*_*i*_) are introduced to simulate a delay [15, 16]. Four sub-states (*h*_1−4_) were empirically chosen as the optimal number for the model to recapitulate experimental data. Modeling the endothelial mechanisms with less than four states resulted in dynamics that did not match experimental data while using more than four states did not notably change the model output. Therefore, when solving CereBRLSIM, eq. 6 is rewritten:

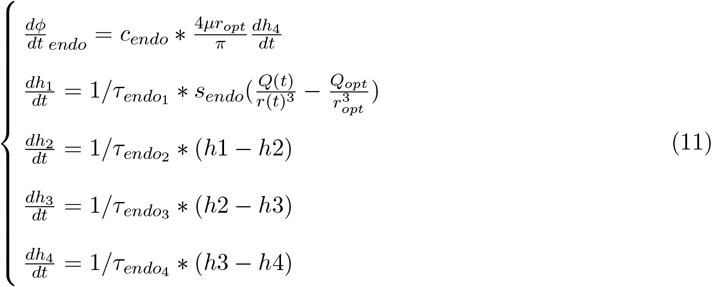

### 7.3 Experimental Model Validation

All dynamic *in vivo* experiments from Fig. 2 were secondary use in this study and were approved by the respective Clinical Research and Ethics Board from the original studies. For each experiment type, the first five patients in the dataset that demonstrated a notable vascular response to intervention were used for model validation.

For each *in vivo* experiment, a large physiological perturbation was performed and the corresponding vascular response was observed. CereBRLSIM was driven with the primary perturbation for each test (pink time courses, Fig. 2). Unless otherwise noted, all other model inputs were assumed to be static. For the pressure-passive scenario, eq. 4 was simulated with 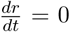 by setting *C*_*myo*_ = *C*_*endo*_ = *C*_*meta*_ = 0. To simulate individual mechanisms only, eq. 4 was solved with *C*_*mech*_ = 1 for the respective mechanism being studied and *C*_*mech*_ = 0 for all other mechanisms.

CereBRLSIM emulates the global cerebral vasculature, whereas experimental measurements correspond to individual arteries where the data were measured. Therefore, we assumed a linear scaling between CereBRLSIM (*x*_*sim*_) and the data (*x*_*data*_): *x*_*sim*_ = *αx*_*data*_ + *β*. To identify this scaling, *α* and *β* were determined for each patient by linear regression between data and full CereBRLSIM simulation (with *α >* 0). This scaling was preserved across all *in silico* experimental conditions for the patient (e.g. pressure-passive model, individual mechanism, etc.).

Cerebral blood velocity (CBv) is CBF divided by the area of the large artery being insonated by Transcranial Doppler. Because large vessels exhibit a smaller change in radius than upstream smaller arterioles[17], we assume that the change in measured radius is 10% of the change in the modeled bulk vascular radius. Therefore, the modeled CBv is calculated as: 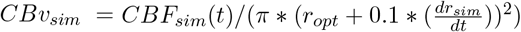.The percentage did not notably impact our findings until it was greater than 50%.

All experiments are described in manuscripts from which they originate. Here, we briefly describe the experiments and the measurements used for model validation.

#### Dynamic cerebral autoregulation (dCA) squat to stand test

Squat to stand protocol was adapted from [18]. After 10 minutes of standing to ensure baseline cardiovascular state, subjects were instructed to alternate between a squatting and a standing position, each held for ten seconds. Beat-to-beat blood pressure was measured by finger photoplethysmography (Finometer Pro, Finapres Medical Systems, Amsterdam, Netherlands) and CBv was measured at the right MCA using 2-MHz transcranial Doppler ultrasound (TCD, Spencer Technologies, Seattle, WA).

#### Flow mediated dilation test

Flow mediation dilation experiments were originally collected in Carr et al.[8], with methodology adopted from [6]. Baseline end-tidal gas concentrations for each subject were established. Participants breathed through an end-tidal forcing system, which provided O_2_ and CO_2_ in proportions appropriate to maintain baseline concentrations. ETCO_2_ was then elevated 9 mmHg above baseline for 30 seconds to invoke a rapid increase in CBv and shear stress. Shear rate time courses, originally calculated in Carr et al.[8], were used in place of 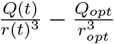 in eq. 11 and simulated diameter was compared to measured diameter of the internal carotid artery (ICA).

#### Neurovascular coupling test

Neurovascular coupling test protocol was adapted from [19, 20]. After two-minutes of resting with eyes closed to ensure baseline vascular and neurologic state, subjects were instructed to open their eyes and read for 30 seconds. ABP was measured with finger photoplethysmography and CBv was measured using 2-MHz TCD at the left posterior cerebral artery (PCA), which has the greatest response in CBv to visual stimulation[20, 21]. Change in brain metabolic state (*ℳ*) was modeled as a step function which increased when eyes opened and decreased when eyes closed.

#### Nonlinear relationships between CBF, ETCO_*2*_, and CPP

To validate that CereBRLSIM could describe the non-linear relationships between change in CBF, ETCO_2_, and CPP, *in silico* experiments were simulated to mimic *in vivo* experiments from Claasen et al.[9] and Klein et al.[17]. In Claasen et al.[9], the relationship between change in CBF and change in ETCO_2_, was assessed by raising or lowering patient ETCO_2_ in approximately 15 seconds. Simulated experiments using the first five patients in table 1 from [9] were conducted with *c*_*myo*_ = 1 and *c*_*endo*_ = *c*_*meta*_ = 10 and *α* = *r*_*opt*_*/*7000. These parameter values were tuned manually to best predict the data. The % change in CBF was calculated as *Q*(*t*)*/Q*_0_. Experimental data in Fig. 3a,c were extracted from Claasen et al. figure 4.

In Klein et al.[17], an open skull piglet model was used to track CBF while ABP was slowly increased or decreased over two hours. *in silico* experiments were conducted by randomly sampling baseline CPP and change in CPP from the distributions reported in table 2 of Klein et al.[17]. A total of 200 *in silico* experiments - 100 increase in CPP and 100 decrease in CPP were conducted. Change in CPP was assumed to be linear. CereBRLSIM was simulated using these approximated patient data with *c*_*myo*_ = 1 and *c*_*endo*_ = *c*_*meta*_ = 0 and *α* = *r*_*opt*_*/*7000. Reported changes in CBF and ABP were calculated by: *CBF*_*t*=2*hrs*_ − *CBF*_*t*=0*hrs*_*/CBF*_*t*=0*hrs*_. Experimental data in Fig. 3d,f were approximated from figure 2c, center panel in Klein et al.[17].

#### 7.4 Traumatic Brain Injury Patient Data

Six two hour patient time series were extracted from the multimodal data set of Traumatic Brain Injury neurocritical care patients at the University of Cincinnati (Table A.2). All patients were male. Patient experiments were chosen to represent the three pressure flow relationships and the full range of patient outcomes. The data utilized in this manuscript include ABP, ICP, PbTO_2_, End Tidal CO_2_, and regional CBF observed by a Bowman probe (https://hemedex.com/products/bowman-perfusion-monitoring-system/). All data were collected under local institutional review board authorization (UCIRB 18-0743).

#### Time series metrics

Mean Flow Index (Mx) and Pressure Reactivity Index (PRx) were calculated according to standard protocol[22]. To calculate Mx, CPP (ABP-ICP) and CBF were averaged over ten second non-overlapping windows. Correlation coefficients between five minutes of averaged samples were calculated using a moving window with 4/5 overlap. These correlation coefficients were then averaged over the one hour dataset. PRx was calculated with the same methodology but correlated ABP and ICP. Pressure Flow Relationships were assessed similar to[23]. First, the linear regression between smoothed CPP and CBF was calculated over one hour of patient data. Time series were labeled positive PFR if the slope was *>* 0.2, zero PFR if slope *<* 0.2 and ≥ -0.1. Negative PFR patients had a slope *<* 0.1.

### 7.5 Assimilating patient data into CereBRLSIM

CereBRLSIM is a forced differential equation: 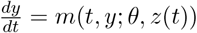, with states *y*, fixed parameters *θ*, and time varying parameters *z*(*t*). All parameter values are given in Table B.1. Patient data are incorporated into the model in two ways: as the input to time varying parameters and used for estimating the CVR parameters.

#### Model inputs

CereBRLSIM time varying parameters are *P, pCO*_2_, *pO*_2_, and 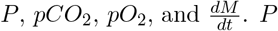 is directly calculated from arteriole blood pressure (ABP) and intracranial pressure (ICP) data. ICP and ABP are each smoothed using a ten-second moving average, down sampled to a resolution of 0.25 Hz, and converted from *mmHg* to *N/cm*^2^ by dividing by 75.0062. *P* is the difference between smoothed ABP and ICP.

Except for cases of ventilation-perfusion mismatch, blood *pCO*_2_ is roughly proportional to the ventilator-derived measurement of end-tidal CO_2_ (*EtCO*_2_). Therefore, *pCO*_2_ in CereBRLSIM is approximated as:

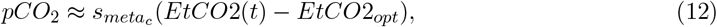

where *EtCO*2_*opt*_ is the 40 mmHg, the nominal value of EtCO_2_ in healthy adults[24].

If arterial oxygenation was measured in our dataset, *PaO*_2_ may be approximated by that input. Instead, *PaO*_2_ is approximated by assuming that oxygen supply is proportional to CBF (that is, *pO*_2_(*t*) ≈ *Q*(*t*)), where *Q* is the simulated *Q* (rather than measured CBF). The ischemic threshold 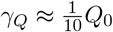 and 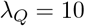.

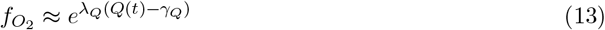

The metabolic activity of the brain (*ℳ*) may ideally be approximated by cerebral metabolic rate of oxygen (CMRO_2_). Rosenthal et al [25] showed a strong relationship between *PbTO*_2_ and *CBF* * (Δ*O*_2_), a subcomponent of *CMRO*_2_. Therefore, while *PbTO*_2_ is not an indicator specific to metabolism, it is correlated with metabolism^1^. Importantly, *PbTO*_2_ is only linearly related to *CBF* * (Δ*O*_2_) below some threshold (*γ*_1_). Therefore, when *PbTO*_2_ data are present, the change in brain metabolic activity is approximated as:

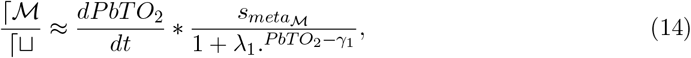

where *λ*_1_ = 10 and *γ*_1_ = 15*mmHg* [25–27] and 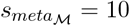.

#### Model Initialization

The magnitude of CBF is dependent on the location of the Bowman Perfusion monitor in the cerebrum. Therefore, a scaling parameter *α* must be estimated that operates as a scaling between *P* and *Q*. In eq. 8 (see also B), length (*L*) is not measurable, but mathematically operates as a scaling between *P* and *Q*. Therefore, we replace *L* with *α* in eq. 8. Before CVR parameters were estimated, *α* was estimated to obtain similar magnitudes of simulated CBF (*Q*) and measured CBF 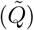.This was done by setting 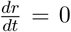 for the first ten minutes of patient data and using interior-point optimization with 200 random starting conditions to solve the optimization problem: 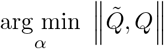.The same estimated *α* was used for pressure passive and CereBRLSIM scenarios.

Cerebral vasculature was assumed to be in resting state at the beginning of the time series. Therefore, *r*_*opt*_, *Q*_*opt*_ and *P*_*opt*_ were set to their initial values (Table B.1). For three patients, setting *P*_*opt*_ = 70 mmHg, the average value of CPP in healthy patients[28] resulted in a lower MSE than using their initial pressure measurement and was chosen instead.

#### Model Personalization

To personalize CereBRLSIM with patient data, we identified the most likely CVR parameter values (*c*_*mech*_ = *{c*_*myo*_, *c*_*endo*_, *c*_*meta*_*}*) given patient CBF data 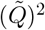^2^. To obtain an uncertainty quantification for the CVR parameter estimate, we assumed a Bayesian framework (eq. 15). We estimated the joint posterior distribution of *{c*_*myo*_, *c*_*endo*_, *c*_*meta*_*}* for each patient given 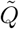 using Markov Chain Monte Carlo Random Walk Metropolis Hastings[30]. Within Gibbs sampling did not notably impact convergence time or results.

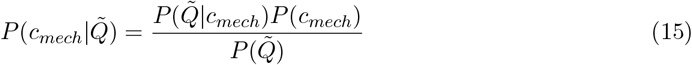

Assuming ergodicity (i.e. multiple chains at the same time is equivalent to one chain for a long time), ten chains were simulated over 100,000 iterations or until convergence was visually confirmed. The last 25% of the chains (or a larger percentage if chains converged quickly) was analyzed. Chain convergence was assessed and chains were removed if either (1) average parameters estimated from a chain resulted in MSE between 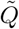 and *Q* greater than half a standard deviation compared to the other chains or (2) removing the chain improved the total Gelman-Reubin statistic. At least two chains were always retained.

### 7.6 CBF Forecasting

After CereBRLSIM was estimated for one hour of patient data, 100 samples were randomly selected from the joint posterior distribution of the CVR parameters. These samples were used to initialize CereBRLSIM, which was then simulated forward for one hour. The sample that minimized the MSE between 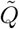 and *Q* was compared with other forecasting approaches.

#### NeuralODE

To compare CereBRLSIM performance against a deep learning approach, neuralODE models were used. NeuralODEs[31] are the most comparable deep-learning approach to CereBRLSIM because they are continuously differentiable and can be simulated forward in time. A feedforward neural network with two fully connected layers with 20 neurons each was trained using the Adam optimizer with 200 mini-batches, each consisting of 200 seconds worth of data. Increased mini-batch size did not impact results. The model was trained for 1200 iterations. Analogous to CereBRLSIM, an individual neuralODE model was trained given CPP time series for each patient using the first hour of patient CBF measurements and then simulated forward for the next hour.

#### Multivariate Linear Regression

To assess how CereBRLSIM compared to a mutlivariate linear regression that leveraged the same datasets, a patient specific multivariate linear regression was trained on the first hour of patient data using inputs of ETCO_2_, CPP, and PbTO_2_ with predictor CBF as the predictor variable. The patient specific multivariate linear regression model was then used to predict CBF for the next hour.

Figures were made in MATLAB, GraphPad PRISM, and Biorender[32].

### 7.7 Data Availability

Experimental data can be accessed from the corresponding authors of the manuscripts from which they originate or by contacting Philip Ainslie. Patient data are available upon request from Brandon Foreman.

## Appendix A Figures and Tables

**Table A.1:**
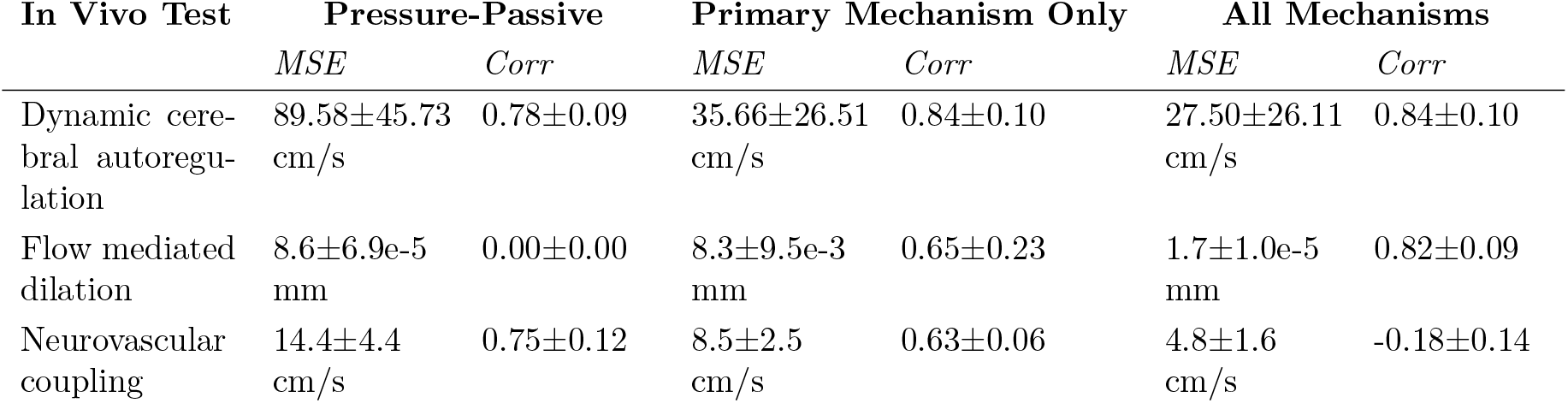
Mean square error and correlation between CereBRLSIM and experimental data, corresponding to Fig. 2.

| In Vivo Test | Pressure-Passive |  | Primary Mechanism Only |  | All Mechanisms |  |
| --- | --- | --- | --- | --- | --- | --- |
|  | *MSE* | *Corr* | *MSE* | *Corr* | *MSE* | *Corr* |
| Dynamic cerebral autoregulation | 89.58±45.73<br>cm/s | 0.78±0.09 | 35.66±26.51<br>cm/s | 0.84±0.10 | 27.50±26.11<br>cm/s | 0.84±0.10 |
| Flow mediated dilation | 8.6±6.9e-5<br>mm | 0.00±0.00 | 8.3±9.5e-3<br>mm | 0.65±0.23 | 1.7±1.0e-5<br>mm | 0.82±0.09 |
| Neurovascular coupling | 14.4±4.4<br>cm/s | 0.75±0.12 | 8.5±2.5<br>cm/s | 0.63±0.06 | 4.8±1.6<br>cm/s | -0.18±0.14 |

**Table A.2:** Neurocritical care patients information. ID corresponds to numerical experimental ID used in manuscript. Experiment ID and Folder corresponds to patient ID and folder within the Cincinnati data set, which experiment starting at start time in hours.

| ID | Experiment ID<br>and Folder | Start<br>Time<br>(hours) | GOSE | PFR | PRx | Mx | $\alpha$ |
| --- | --- | --- | --- | --- | --- | --- | --- |
| 1 | 1074 02 | 18.7 | 1 | p | -0.14 | 0.85 | 49.9 |
| 2 | 1086 01 | 16.1 | 1 | p | 0.06 | 0.86 | 40.0 |
| 3 | 1185 01 | 3.7 | 8 | z | 0.28 | 0.17 | 2.2 |
| 4 | 1184 01 | 9.2 | 4 | z | 0.06 | 0.11 | 15.0 |
| 5 | 1166 04 | 11 | 3 | n | 0.51 | 0.34 | 10.5 |
| 6 | 1184 02 | 25.2 | 4 | n | 0.48 | 0.35 | 3.3 |

**Table A.3:**
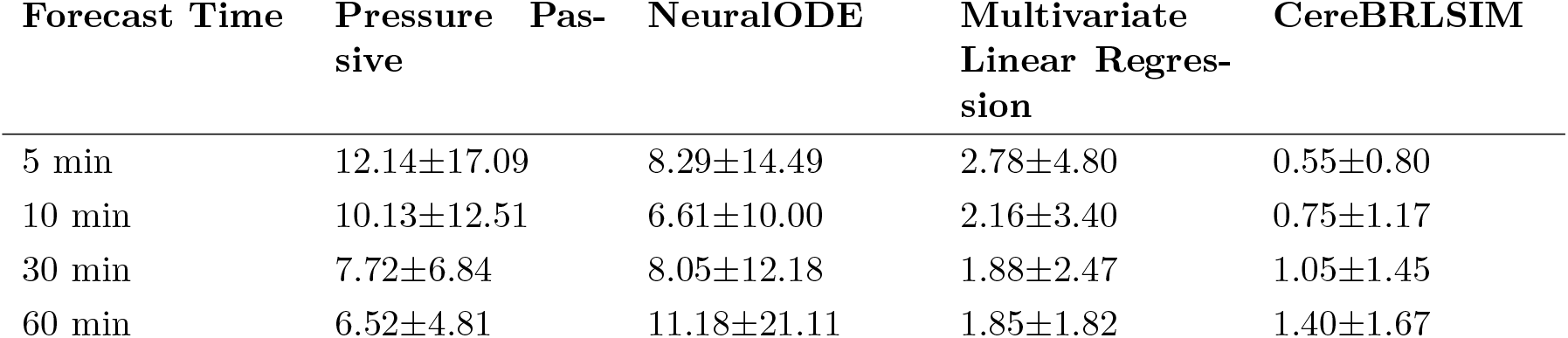
Raw MSE values for simulating CBF forward 5, 10, 30, and 60 minutes into the future after personalization or model training.

**Figure A.1:**
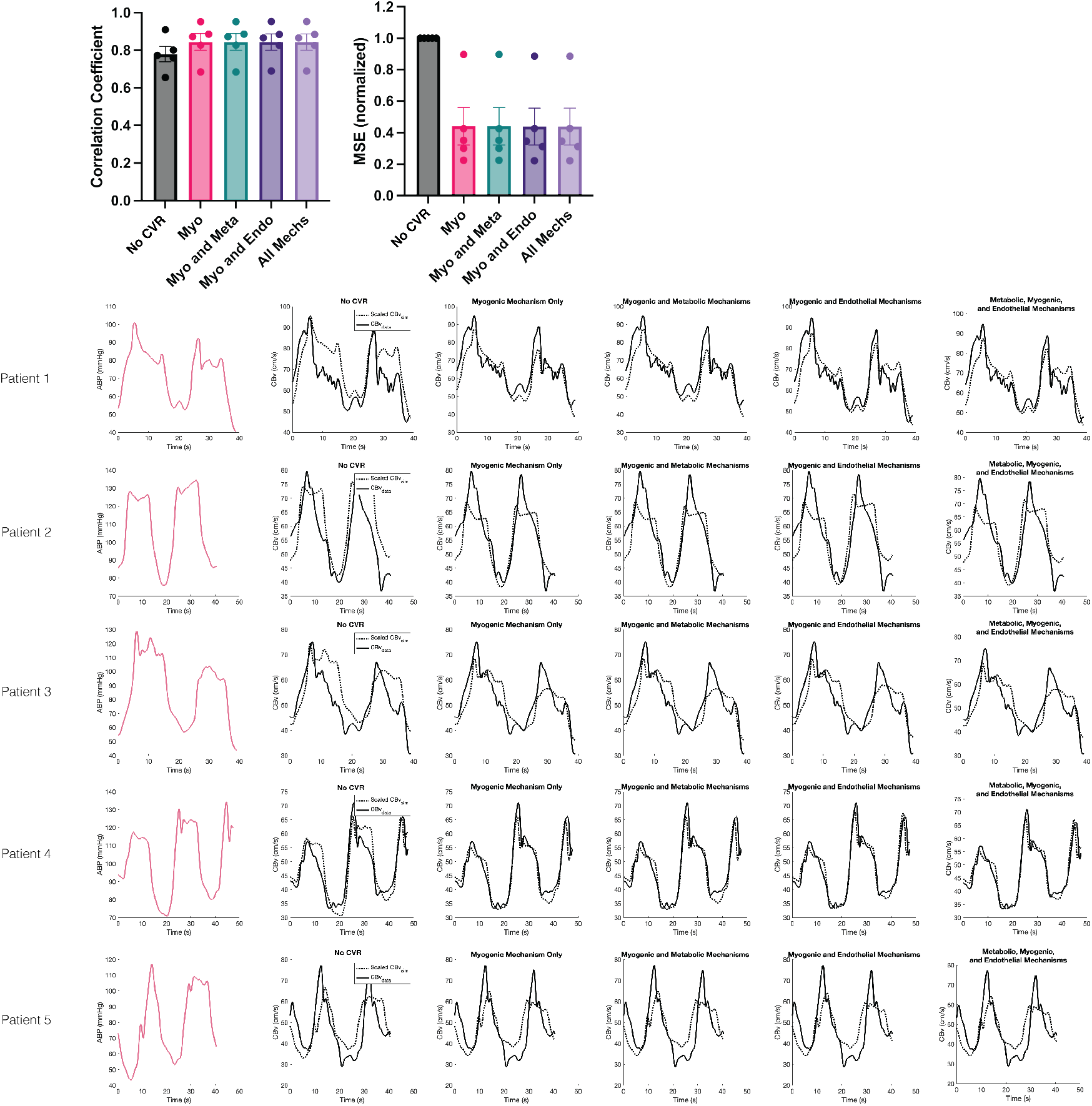
Time courses for all patients and all CereBRLSIM experiments for squat to stand test.

**Figure A.2:**
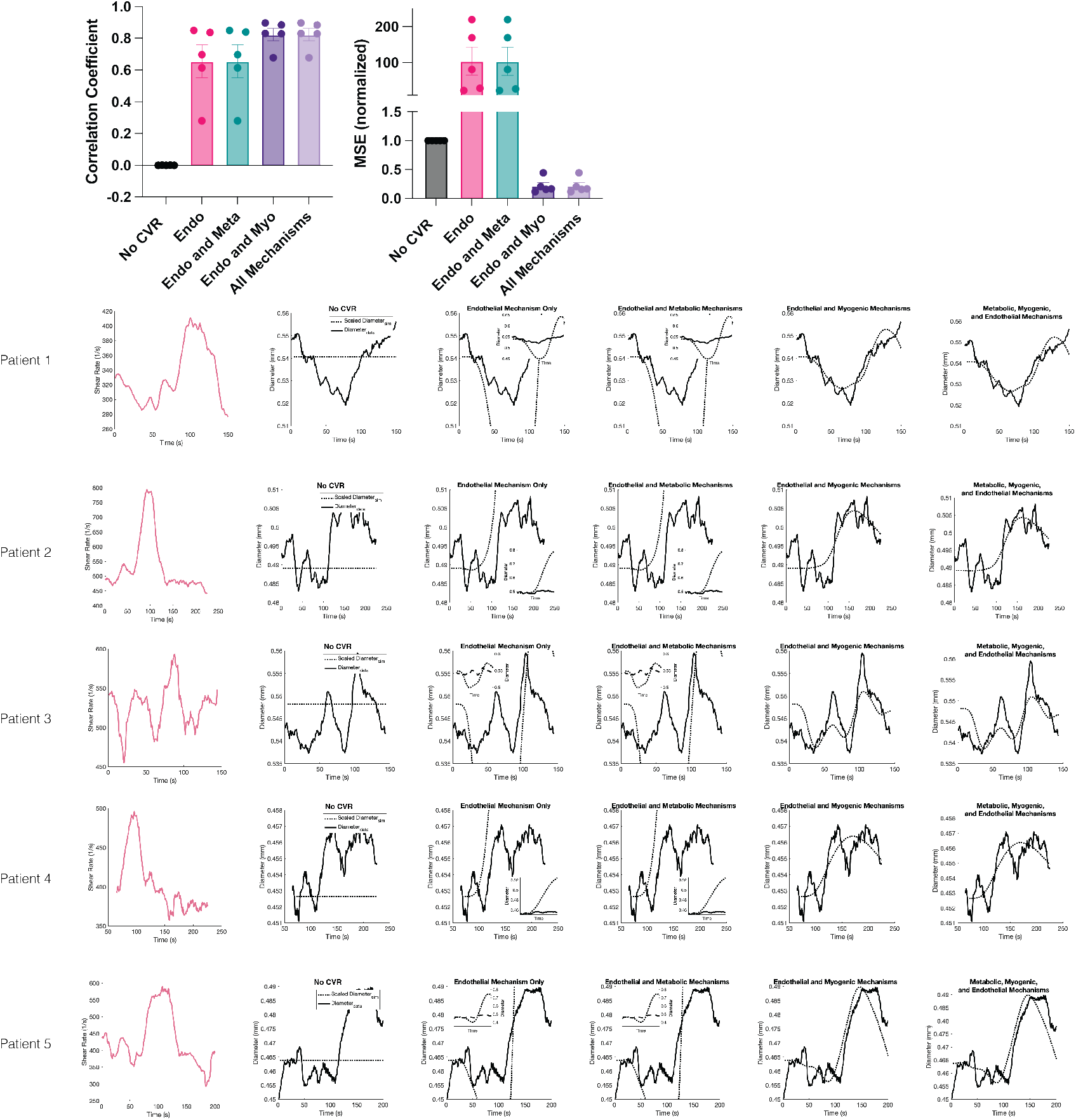
Time courses for all patients and all CereBRLSIM experiments for flow mediated dilation test.

**Figure A.3:**
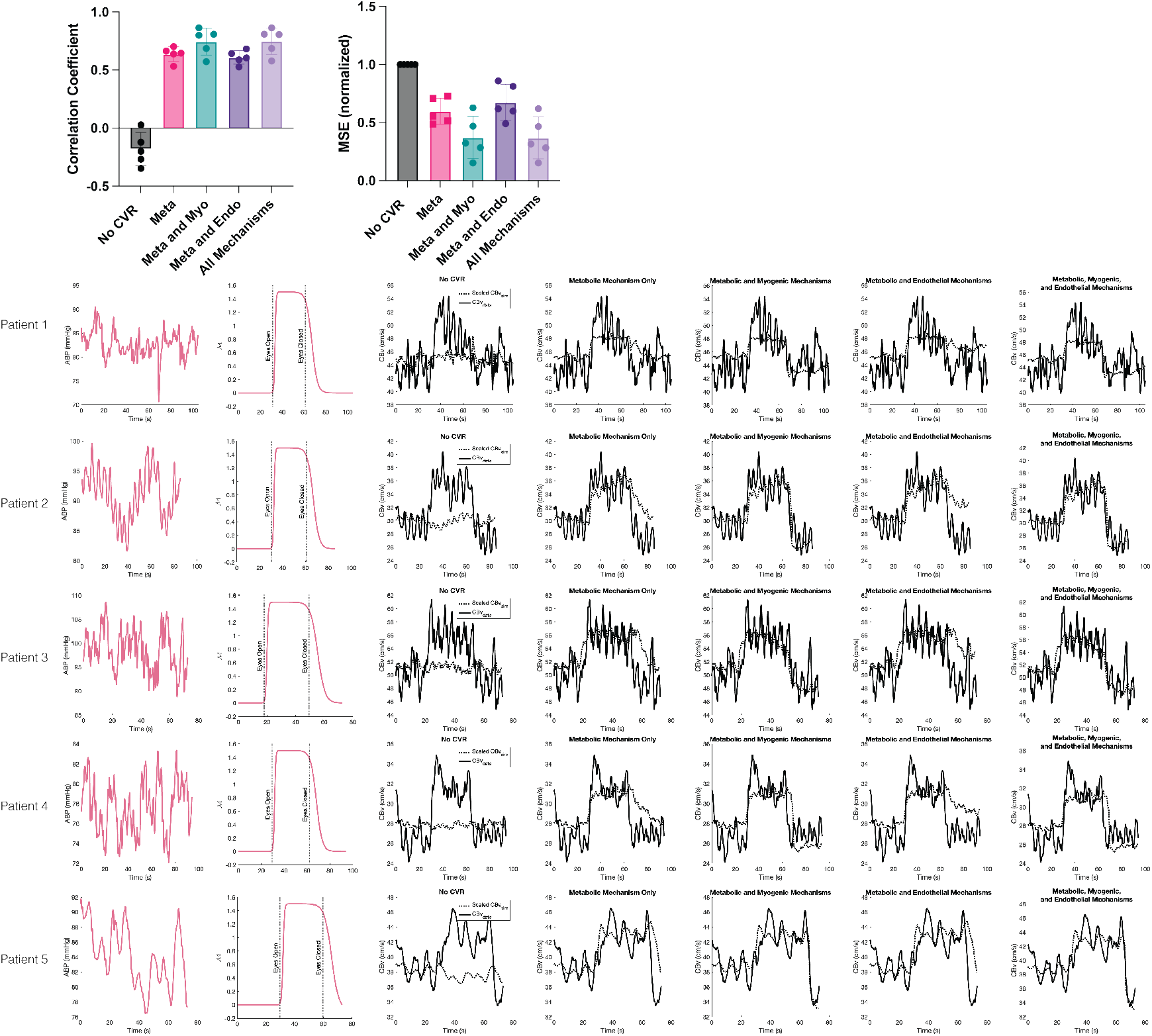
Time courses for all patients and all CereBRLSIM experiments for neurovascular coupling test.

**Figure A.4:**
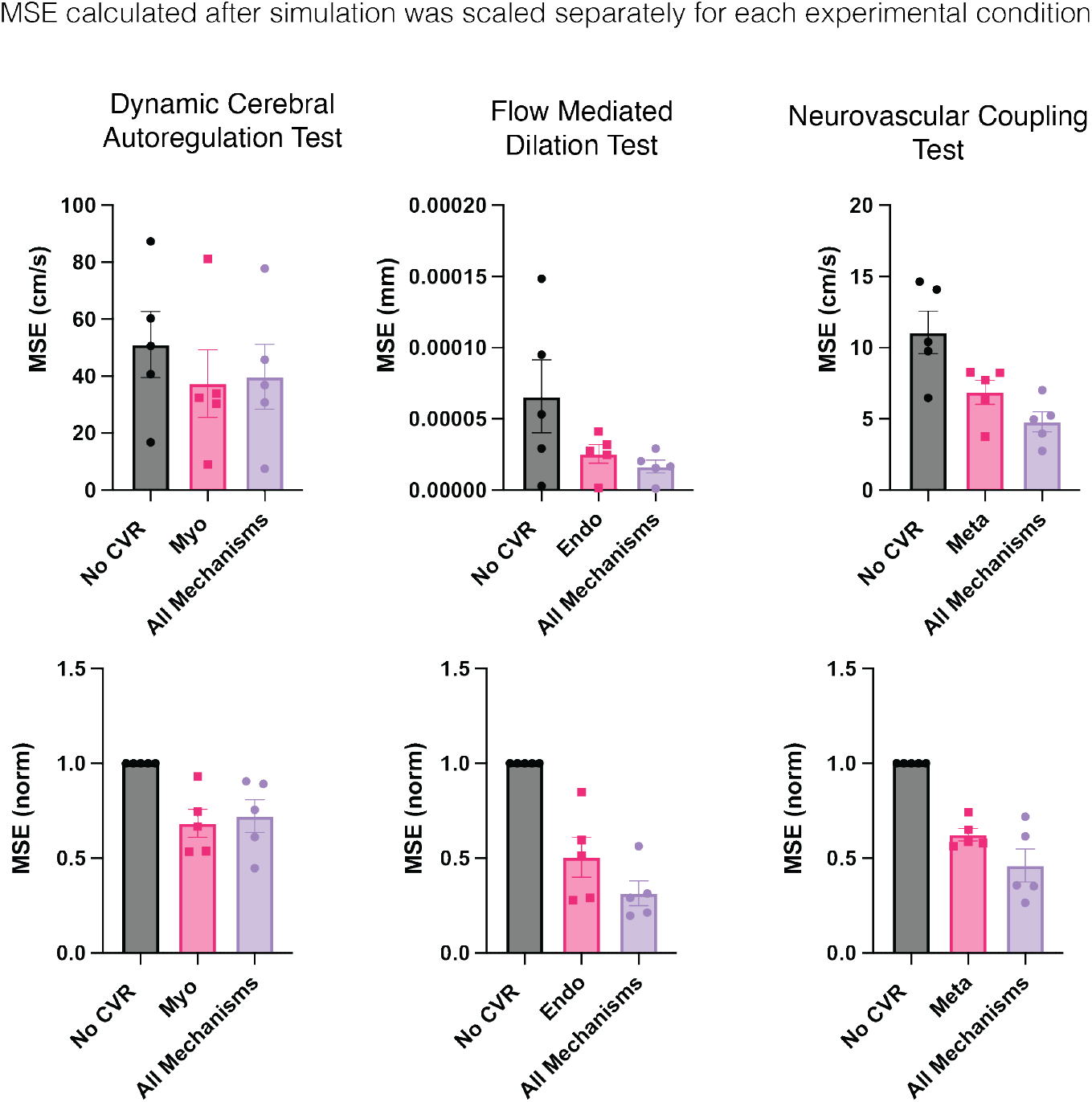
Mean square error between model prediction and data when each model condition was scaled individually.

**Figure A.5:**
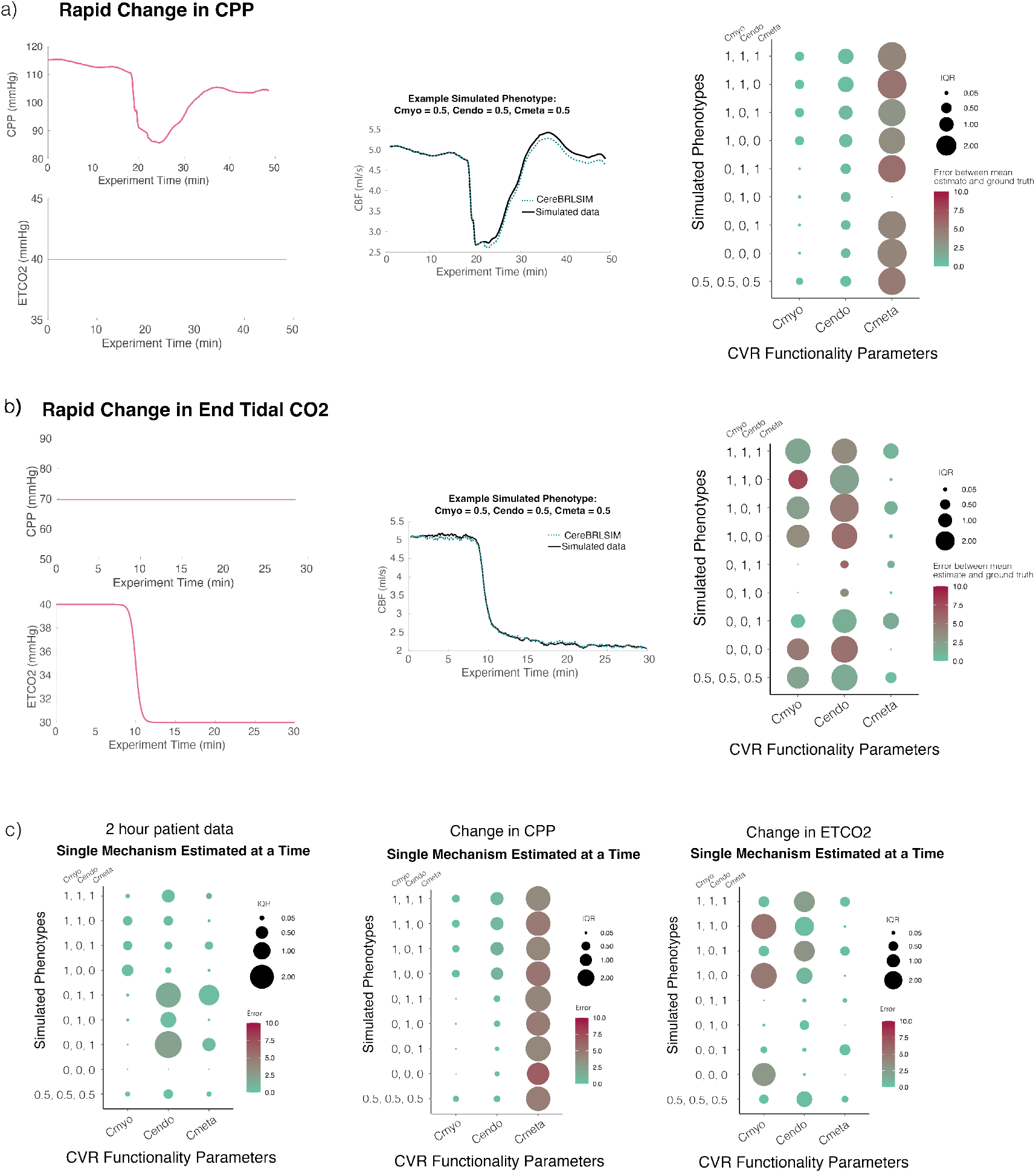
Additional CVR Parameter Estimatimability Experiments. (a) Experiment assessing parameter estimability when CPP changes but ETCO2 is held fixed. (b) Experiment assessing parameter estimability when ETCO2 changes and CPP is held fixed. (c) Results of estimating individual parameters while other two are set to their correct value. Left plot corresponds to Fig. 4, middle plot corresponds to (a), right plot corresponds to (b).

**Figure A.6:**
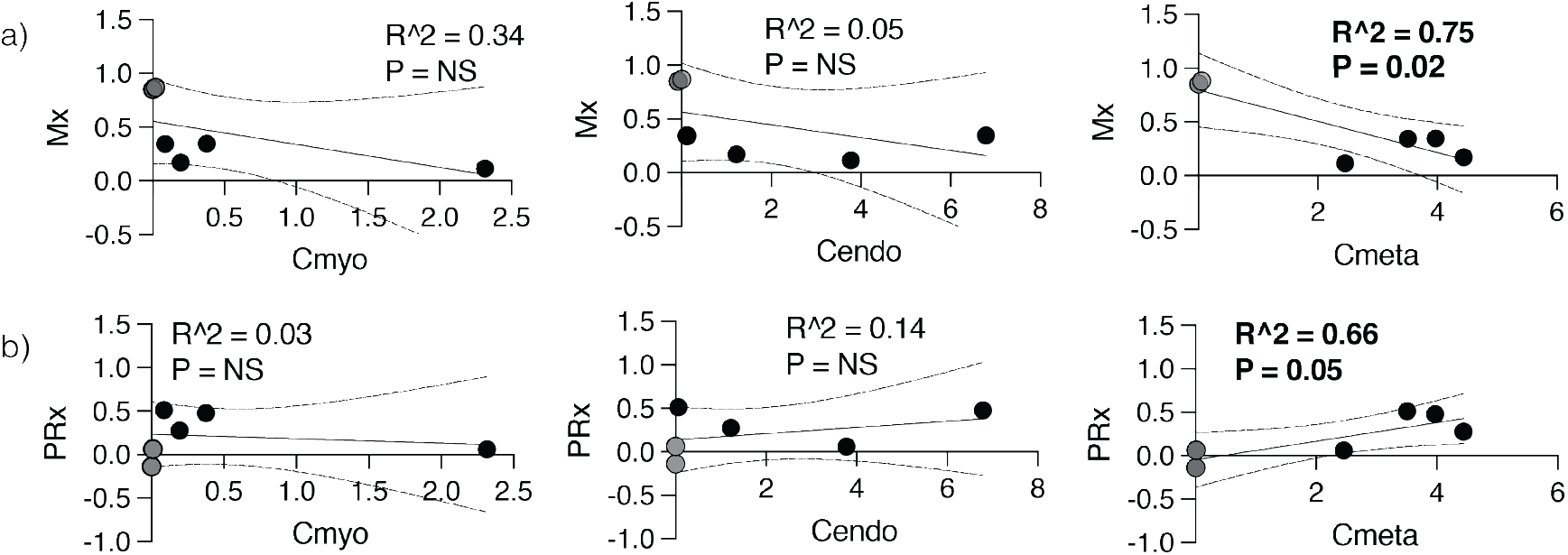
Correlation between CVR parameters and other metrics of vascular function. Patients with positive pressure flow relationships are shown in gray. **(a)** Mean flow index (Mx) is a correlation coefficient between CPP and flow and is a proxy for cerebral autoregulation, the pressure-dependent subcomponent of CVR. **(b)** Pressure reaCctivity index (PRx) is a correlation coefficient between ICP and ABP and is commonly used as a proxy for cerebral autoregulation in the absence of CBF metrics. Colors and numbers of each patient correspond to patients in Fig. 5.

## Appendix B Equations

The full CereBRLSIM model as it is coded for integration with neurocritical care patient data is given here:

Change over time is denoted as 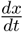.External model drivers are denoted with bold symbol (***x***). Note that length (*L*) is replaced with scaling parameter *α* (Supp. C.6).

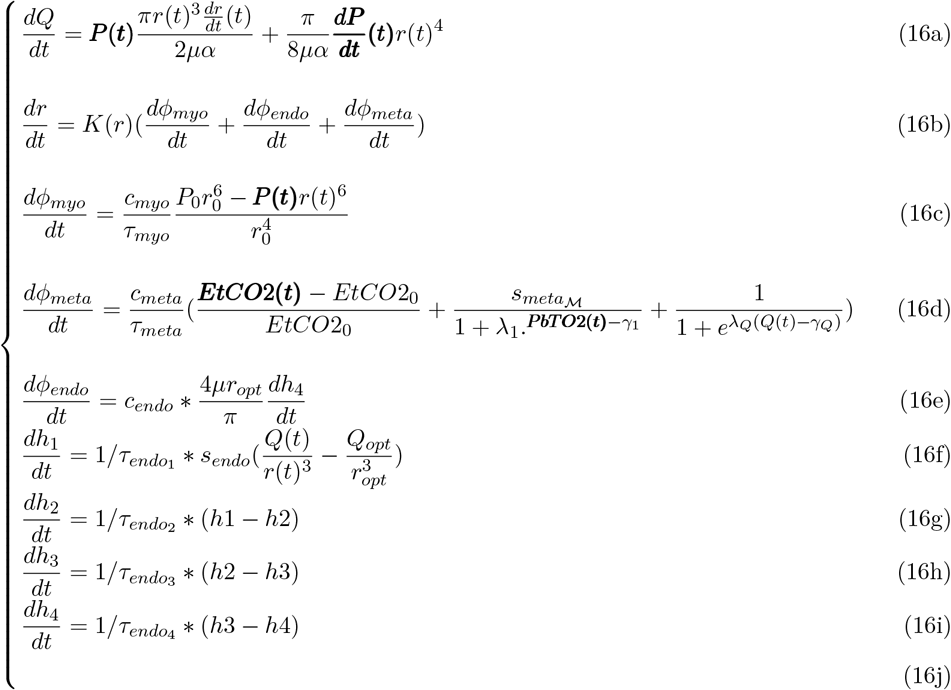

*K* represents the length tension relationship in the vasculature given by:

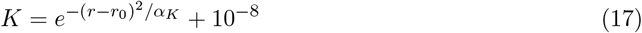

**Table B.1:**
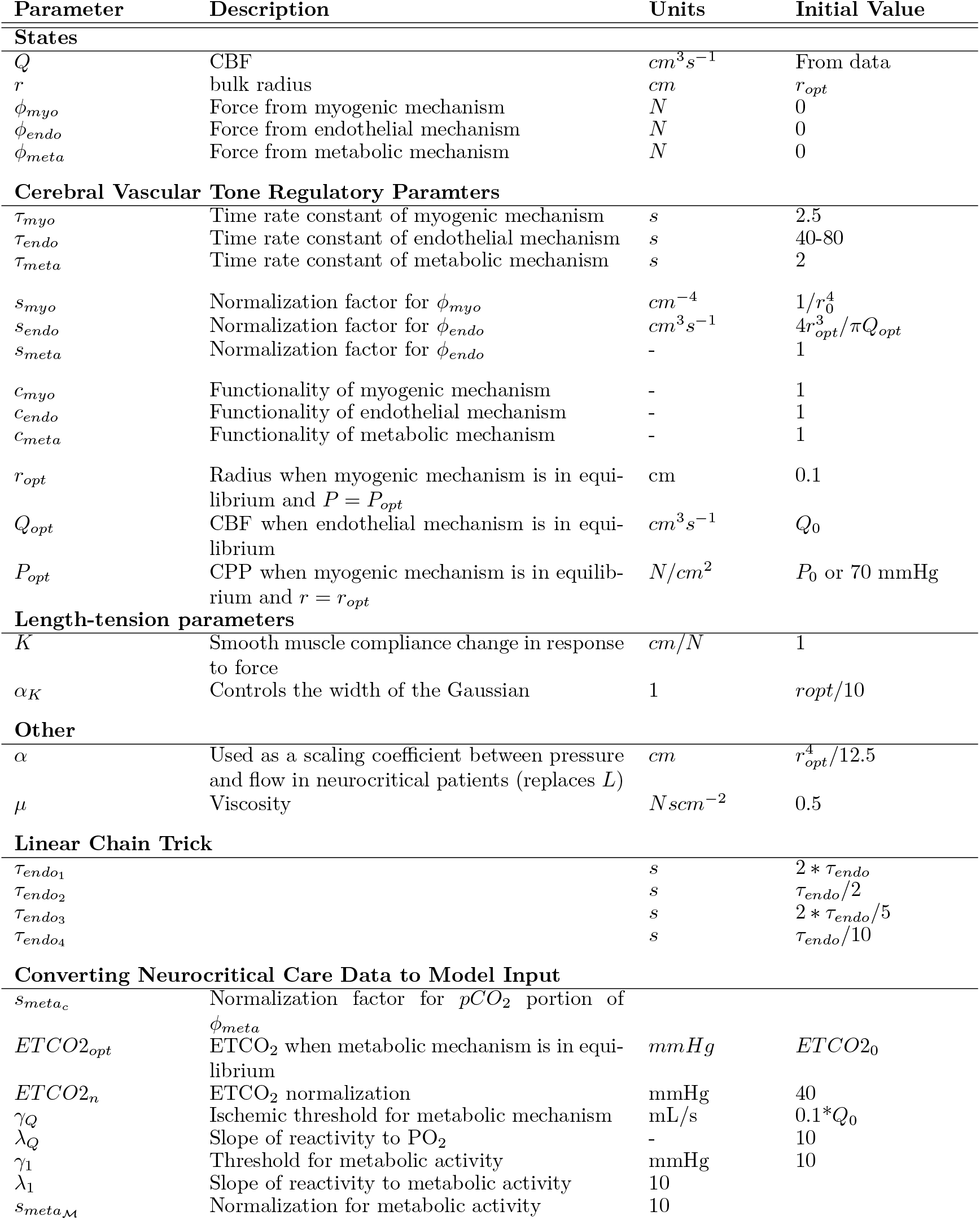
All relevant parameters with units and values. - indicates no units. Unless otherwise stated, initial values, indicated as *X*_0_ are calculated as the first value in the patient time series.

| Parameter | Description | Units | Initial Value |
| --- | --- | --- | --- |
| **States** |  |  |  |
| $Q$ | CBF | $cm^3 s^{-1}$ | From data |
| $r$ | bulk radius | $cm$ | $r_{opt}$ |
| $\phi_{myo}$ | Force from myogenic mechanism | $N$ | 0 |
| $\phi_{endo}$ | Force from endothelial mechanism | $N$ | 0 |
| $\phi_{meta}$ | Force from metabolic mechanism | $N$ | 0 |
| **Cerebral Vascular Tone Regulatory Paramters** |  |  |  |
| $\tau_{myo}$ | Time rate constant of myogenic mechanism | $s$ | 2.5 |
| $\tau_{endo}$ | Time rate constant of endothelial mechanism | $s$ | 40-80 |
| $\tau_{meta}$ | Time rate constant of metabolic mechanism | $s$ | 2 |
| $s_{myo}$ | Normalization factor for $\phi_{myo}$ | $cm^{-4}$ | $1/r_0^4$ |
| $s_{endo}$ | Normalization factor for $\phi_{endo}$ | $cm^3 s^{-1}$ | $4r_{opt}^3/\pi Q_{opt}$ |
| $s_{meta}$ | Normalization factor for $\phi_{endo}$ | - | 1 |
| $c_{myo}$ | Functionality of myogenic mechanism | - | 1 |
| $c_{endo}$ | Functionality of endothelial mechanism | - | 1 |
| $c_{meta}$ | Functionality of metabolic mechanism | - | 1 |
| $r_{opt}$ | Radius when myogenic mechanism is in equilibrium and $P = P_{opt}$ | cm | 0.1 |
| $Q_{opt}$ | CBF when endothelial mechanism is in equilibrium | $cm^3 s^{-1}$ | $Q_0$ |
| $P_{opt}$ | CPP when myogenic mechanism is in equilibrium and $r = r_{opt}$ | $N/cm^2$ | $P_0$ or 70 mmHg |
| **Length-tension parameters** |  |  |  |
| $K$ | Smooth muscle compliance change in response to force | $cm/N$ | 1 |
| $\alpha_K$ | Controls the width of the Gaussian | 1 | $r_{opt}/10$ |
| **Other** |  |  |  |
| $\alpha$ | Used as a scaling coefficient between pressure and flow in neurocritical patients (replaces $L$) | $cm$ | $r_{opt}^4/12.5$ |
| $\mu$ | Viscosity | $N scm^{-2}$ | 0.5 |
| **Linear Chain Trick** |  |  |  |
| $\tau_{endo1}$ | | $s$ | $2 * \tau_{endo}$ |
| $\tau_{endo2}$ | | $s$ | $\tau_{endo}/2$ |
| $\tau_{endo3}$ | | $s$ | $2 * \tau_{endo}/5$ |
| $\tau_{endo4}$ | | $s$ | $\tau_{endo}/10$ |
| **Converting Neurocritical Care Data to Model Input** |  |  |  |
| $s_{meta_c}$ | Normalization factor for $pCO_2$ portion of $\phi_{meta}$ | | |
| $ETCO2_{opt}$ | $ETCO_2$ when metabolic mechanism is in equilibrium | $mmHg$ | $ETCO2_0$ |
| $ETCO2_n$ | $ETCO_2$ normalization | mmHg | 40 |
| $\gamma_Q$ | Ischemic threshold for metabolic mechanism | mL/s | $0.1 * Q_0$ |
| $\lambda_Q$ | Slope of reactivity to $PO_2$ | - | 10 |
| $\gamma_1$ | Threshold for metabolic activity | mmHg | 10 |
| $\lambda_1$ | Slope of reactivity to metabolic activity | 10 | |
| $s_{meta_M}$ | Normalization for metabolic activity | 10 | |

## Appendix C Mathematical Justification of Study Assumptions

The overall goal of this study is to build a digital twin that can predict cerebral blood flow (CBF) using time series data available in neurocritical care. There are numerous ways to build such a digital twin. This section provides a detailed rationale for the assumptions and decisions made in our study. Each assumption introduces its own limitations, inevitably creating a divergence between our model and the real-world system it represents. However, the assumptions we adopt are well-established in the field and, in our opinion, are justified given both the objectives of this study and the limited data available.

We begin by describing the foundational fluid dynamics models that our models are motivated by (Section C.1), including the assumptions made for their construction. The next section (Section C.2) briefly describes physiology that we believe must be addressed before solving applying these models to cerebral vascular domain. The rest of the section addresses this physiology by explicitly stating assumptions we make in the modeling process and describing specific cases and scales on which these assumptions (and therefore our model) can be justified. Throughout the section, we highlight all relevant assumptions with the nomenclature:

### Assumption 0.

*State assumption here*.

Assumptions that we believe are particularly important in our context are further highlighted by outline. Many of the following descriptions are motivated by an excellent text book by Zamir [1].

### C.1 Foundation Models of Fluid Dynamics

We aim to develop a parsimonious model to represent CBF using continuously available time series data from neurocritical care. To achieve this, we first examine two foundational models from fluid dynamics to determine what modifications are needed based on our system and available data.

#### C.1.1 Navier-Stokes

The term ‘flow’ refers to the change in position of fluid over time. The governing equations used to describe fluid flow in three spatial dimensions (**v**) are the Navier-Stokes equations.

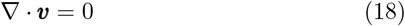

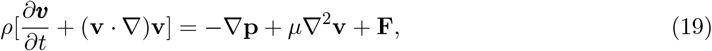

The first equation (eq. 18) states that mass is conserved in the system. The second equation (eq. 19) states that momentum is conserved and is based on Newton’s second law: *F* = *ma*. Here, the right hand side of the equation gives the force terms. The term ∇**p** is the pressure (*p*) acting on the fluid in all directions. The term *µ*∇^2^**v** represents resistance to flow due to viscous interactions, and the term **F** describes external forces per unit volume. The left hand side of the equations are the inertial or ‘*ma*’ terms, where *ρ* is the density of the fluid (*ρ* = *m/V*). Implicit in the Navier-Stokes equations are two key assumptions that are relevant in our application to cerebral vascular dynamics:

##### Assumption 1.

*The fluid is incompressible and moves in a roughly Newtonian manner*

##### Assumption 2.

*Mass is conserved*^3^

Practical implementation of the Navier-Stokes equations requires imposing a domain through which the fluid flows. It is helpful to start with a simplistic domain, such as a single straight tube. Flow through a tube is described by Poiseuille’s law.

#### C.1.2 Poiseuille

Poiseuille was a physician and experimentalist who studied blood flow through frog capillaries[2]. He empirically constructed an equation for blood flow through a single vessel as:

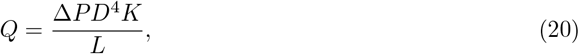

where *L* is length of the tube, *D* is the diameter of the tube, and *K* is a constant parameter dependent on temperature and the type of fluid. Δ*P* is the gradient of pressure along the axis of the tube. *Q* is average volumetric flow. This equation is not what is typically thought when considering “Poiseuille’s law”. Rather, Hagenbach (and likely Stokes) derived eq. 21 and named it “Poiseuille’s law” because the functional form closely matched eq. 20.

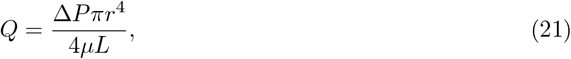

where *r* is the vessel radius and *µ* is the viscosity.

**Figure C.1:**
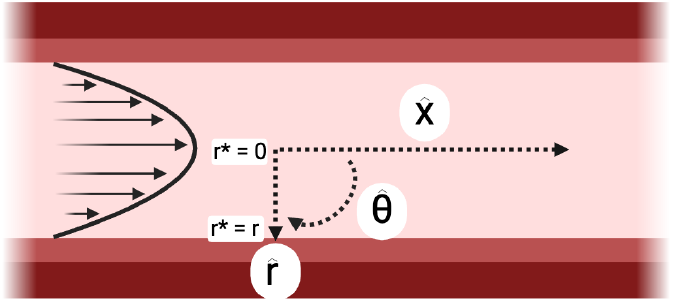
Poiseuille flow through a tube takes a parabolic shape, with largest velocity at the center of the tube (*r*^*^ = 0). Cylindrical coordinates for the tube are given to the right.

Eq. 21 can be derived from the Navier-Stokes equations by writing eqs. 18 and 19 in cylindrical coordinates (Fig. C.1) making the following assumptions:

##### Assumption 3.

*pressure and flow in the radial* 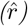 *or circumferential* 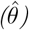 *directions are negligible compared to the axial direction* 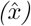

##### Assumption 4.

*External forces are negligible (F* = 0) ^4^,

leading to the formulae:

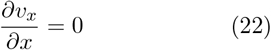

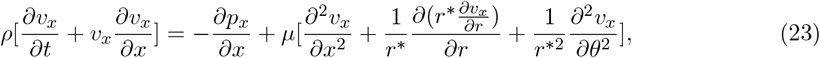

where *r*^*^ is the radial distance of each particle from the center of the tube, not to be confused with the radius of the vessel (*r*).

Assuming:

##### Assumption 5.

*second order terms are negligible*,

and substituting the conservation of mass equation (eq. 22) into eq. 23 gives:

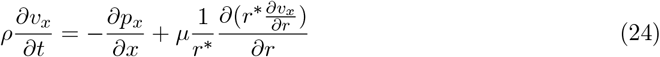

From eq. 24, we observe that there are two terms operating on velocity: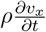 and 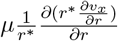. The first term 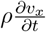, is the inertial term operating in the *x* direction. Inertia, is the concept commonly described by the colloquialism: “an object in motion stays in motion unless acted upon by an outside force” (or equivalently, an object at rest stays at rest). Therefore, inertia can also be thought of as an intrinsic “resistance” to changes in flow. The second term 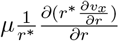,is the viscous term, which describes the resistance to flow due to viscous interactions. Depending on the system, the flow behavior may be categorized in three ways: inertial effects dominate (turbulent flow), viscous effects dominate (laminar flow), or effects are both present (transitional flow). The characteristic flow profile can be predicted using quantities like the Reynolds number (for steady flow profiles) or the Womersley number (for pulsatile flow profiles).

To derive Poiseuille flow, assume:

###### Assumption 6.

*Steady State Assumption: flow is dominated by viscous forces:* 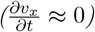.

This assumption allows simplification of eq. 24:

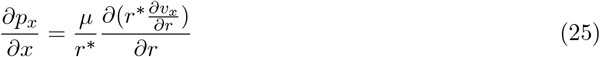

Integrating over the length of the vessel: 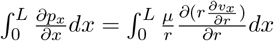, gives:

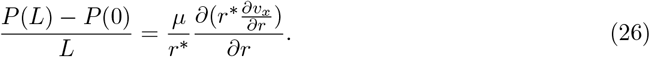

Defining

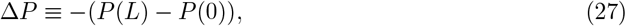

and substituting eq. 27 into eq. 28 gives:

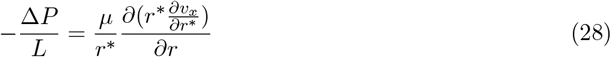

##### Remark 1.

Integration takes the continuous average over space or time, trading the ability to resolve ‘fine resolution’ quantities (up to the integration scale) for average or bulk quantities. For example, here, integration removes mathematical ability to track differences in blood flow across the axis of the tube but provides an equation for average flow along the tube.

To obtain eq. 28 in terms of *v*_*x*_, we integrate both sides along the radial direction.

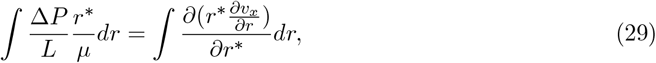

leading to the equation:

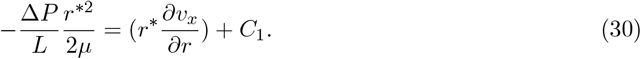

Rearranging and integrating again over r yields:

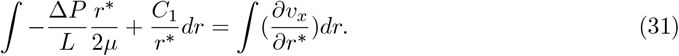

Evaluating over the integral results in:

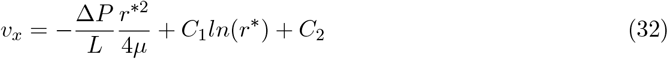

To obtain values for the constant terms, we make the following assumptions:

##### Assumption 7.

*Velocity of flow at the center of the tube (r*^*^ = 0*) is finite*.

This assumption requires *C*_1_ = 0. Also assume the

##### Assumption 8.

*no slip condition: velocity at the walls is zero, v*_*x*_(*r*^*^ = *r*) = 0.

This assumption leads to the equation for *C*_2_:

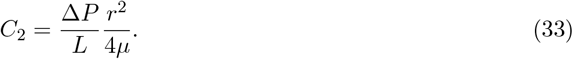

Therefore, the final equation for *v*_*x*_ is given by:

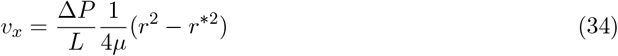

It can be seen from this equation that viscous forces cause the fluid to form a parabolic (Fig. C.1), in which the maximum velocity 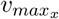 occurs at the center of the tube *r*^*^ = 0:

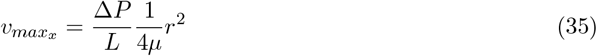

To obtain the average flow, integrate over the circumference divided by the total tube area:

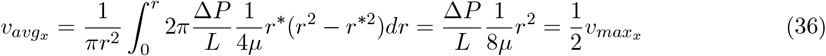

Therefore, the average volumetric flow is *Q* = 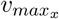 * *πr*^2^ which is fully described by
substituting eq. 35 into eq. 36 and multiply by area:

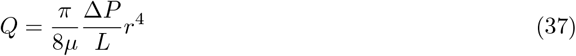

Equation 37 is canonically known as Poiseuille’s law or the Hagen-Poiseuille equation.

### C.2 Foundational Understanding of Cerebral Vascular Physiology

The Navier-Stokes and Poiseuille equations give us a starting place for modeling fluid motion through a system. To apply these equations, domains and boundary conditions must be imposed upon them, meaning we must have an understanding of what the fluid is flowing *through* and what the forces on the fluid are. In this section, we discuss specifics of the cerebral vascular system that we believe should be considered.

#### C.2.1 Complex Vascular Tree

The vascular system is a network of blood vessels that extend throughout the body to transport oxygen, nutrients, and waste to and from the organs. The heart is a physiological pump that creates strong pressure oscillations to fill with blood from veins, facilitate blood re-oxygenation via the lungs, and push blood back out through the body via arteries. Nearly 20% of the blood leaving the heart is directed into the cerebral vasculature. The cerebral vasculature is a highly complex network of branching vessels.

This branching network is important for fluid flow for multiple reasons. Specifically, when flow meets a point in space where the vessels branch, known as a bifurcation, some fluid may be reflected backwards. The impact of these bifurcations becomes more complex when pressure is dynamic, which is the case for the vascular system.

#### C.2.2 Semi-pulsatile Dynamic Pressure Gradient

The heart rhythmically contracts and relaxes in a semi-pulsatile manner to produce large pressure pulses that accelerate blood through arteries. As such:

##### Assumption 9.

*blood flow throughout the body, including cerebral blood flow, is primarily pressure driven (as opposed to being primarily driven by turbulent or viscous effects)*.

The pressure created by the heart is called arterial blood pressure (ABP). The pulsatile nature of ABP has implications for the flow profile. One of those implications is that it impacts whether flow is dominated by inertial or viscous forces. Another is that, because vessels are compliant (discussion to follow), pulsatile pressure induces traveling waves where the vessels ‘bulge’ and then travel along the tube axis.

In addition to pulsatile forcing pressure, one thing that makes cerebral hemodynamics unique is the pressure at the end of the arteries (*P* (*L*)). The skull is a hard object. This means that there is an upper bound for the total amount of volume available for materials including the brain and blood vessels to occupy. Pressure in the skull created because of its fixed volume is called the intracranial pressure. Generally, pressure at the end of the arteries is thought to be equivalent to intracranial pressure because of the following argument.

The cerebral vascular system comprises of arterioles bringing blood and nutrients to the brain, which diverge into thin walled capillaries through which the nutrients diffuse out of the vessels, which subsequently converge into capillaries. The study of blood flow is usually concerned with how the brain is perfused (however, the ability of blood to leave the cerebral veins is also important to study). Therefore, we aim to model blood flow through the arterioles.

Recall eq. 27, which defined Δ*P* as the pressure difference between the tube entrance (*P* (0)) and tube exit (*P* (*L*)). Temporarily assuming that blood flows from the heart through the brain in a single artery (this assumption is discussed in depth in section C.4), then *P* (0) is ABP and *P* (*L*) is the pressure at the start of the capillaries and veins. As opposed to arterioles (discussed later), capillaries and veins are highly compliant and cannot form substantial wall tension (assuming the pressure is not higher than the vessel’s yield strength, or the point at which vessel deformation is no longer temporary.) Laplace’s law for cylinders states that wall tension *T* is given by:

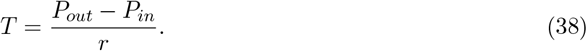

Because *T* ≈ 0 in capillaries and veins,

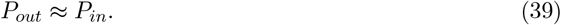

Therefore, the pressure at the inlet of the capillaries and veins, and subsequently, the outlet of the arterioles: *P* (*L*) ≈ ICP. The pressure gradient driving blood through the cerebral vascular arteries, is generally thought to be:

##### Assumption 10.

*Inlet pressure (P* (0)*) is approximated by arterial blood pressure (ABP) and outlet pressure (P* (*L*)*) is approximated by ICP*.

This assumption is invalid if something is blocking venous outflow, causing *P*_*in*_ to become larger than *P*_*out*_.

#### C.2.3 Dynamic Radius

Cerebral arteries and arterioles are not rigid tubes. Instead, their radius can change both “passively” and “actively”. In vivo, vessels are constantly subjected to transmural pressure (*P*_*T*_), defined as the difference between the pressure inside and outside the vessel. For the cerebral vasculature, internal pressure is proportional to ABP, while external pressure is governed by ICP. Cylindrical vessel wall *T* is given by Laplace’s law: *T* = *P*_*T*_ *r*. If the vessel cannot create tension to oppose changes in *P*_*T*_, the radius will passively change size (as discussed for the capillaries and veins).

Different from most capillaries and veins, arteries and arterioles are surrounded by varying degrees of smooth muscle, which can produce tension. Smooth muscles are made up of layered actin and myosin filaments. The degree of overlap controls the vessel radius. To control this overlap, small functional units, known as cross-bridges, extend from the myosin filament and form attachments with the actin filament. The formation of these cross-bridge attachments, also known as *cross-bridge activation*, is constantly controlled by chemical processes.

Activated cross-bridges create tension to oppose transmural pressure from passively changing vessel radius. When radius is not changing, but cross-bridges are exerting force to oppose transmural pressure, this opposing force is known as isometric contraction. Isometric contraction force is modulated the number of activated cross-bridges (explained nicely in [3]). Even in resting state, smooth muscle cross-bridges are constantly cylcing (detaching and reattaching) to maintain a basal radius slightly smaller than *r*_0_. The amount of energy consumed by cross-bridges is determined in part by how fast they cycle. Smooth muscle cross bridges cycle much slower than skeletal muscles. Slow cycling cross-bridges are sometimes called “latch-bridges”[4].

Despite smooth muscle isometric contraction, arteries and arterioles do exhibit some passive changes in radius. The extent that the vessels change volume (and proportionally radius) for a given in pressure (*dV/dP*) is called the vascular compliance *C*. Activated cross-bridges can act as linear springs, allowing slight stretch under changes in transmural pressure. However, the contribution of cross-bridges to passive radius change is thought to be relatively small compared to contributions by elastin and other elastic fibers that make up the vessel[3]. Therefore, compliance of arterioles is primarily influenced by proportion of elastin to smooth muscle comprising the vessel wall. Cerebral arteries have less elastin[5] and are generally less compliant[6] than systemic arteries.

In addition to maintaining vessel radius via isometric contraction, an important role of the smooth muscle is its ability to actively change vessel radius. After activation, cross-bridges may undergo another conformational changed called a ‘power-stroke’. During a power-stroke, the myosin head ‘bends’ over, sliding the actin and myosin filaments over each other to increase overlap. Successive power-strokes by cross bridges can result in large decreases in vessel radius. This is known as concentric contraction. Alternatively, cross-bridges can also allow for ‘active dilation’. As previously discussed, transmural pressure is typically positive and constantly exerting force on the vessel to dilate. If all cross-bridges were de-activated (detached from actin), the vessel would passively dilate proportional to pressure and vessel compliance. Alternatively, cross-bridges may facilitate dilation actively by gradually reducing the number of activated cross-bridges while maintaining activation in others, allowing transmural pressure to increase vessel radius in a controlled manner. This is called eccentric contraction. The contractile state of the cross-bridges dictates the cerebral vascular tone.

Eccentric and concentric contraction are both initiated by complex biomechanical triggers. These triggers are grouped into three categories: myogenic, metabolic, and endothelial, which we collectively refer to as cerebral vascular regulation (CVR). The myogenic mechanism triggers changes in radius to compensate for changes in pressure. The metabolic mechanism triggers changes in radius to compensate for changes in metabolic demand. The endothelial mechanism triggers changes in radius in response to shear stress. These triggers are described more thoroughly in the manuscript text.

Mechanisms that actively maintain or change vessel radius can become dysfunctional. We refer to this functionality as “active vessel function”. Functionality of CVR mechanisms (which dictate active changes in vessel radius) are a primary focus of the main manuscript.

#### C.2.4 Summary

We aim to model continuous CBF. There are three primary characteristics of the cerebral vascular system which we believe are particularly important to consider for this model:

1. Cerebral vasculature has a highly complex tree-like structure
2. Driving pressure is pulsatile
3. Vessel radius can change both actively and passively

In the following sections, we explore how the fluid dynamic models described in section C.1 can be applied given these physiological considerations.

### C.3 Justifying Steady State, Laminar Poiseuille Flow

Poiseuille’s law assumes that flow is in steady state and remains smooth or laminar (assumption 6). The steady flow assumption is justifiable if viscous effects dominate inertial forces to suppress the development of turbulence, which is quantified by evaluating their ratio. Two common calculations for this ratio are Reynolds number and Womersley’s number.

In human vasculature, the Reynolds number decreases with distance from the heart. Reynolds number is approximately 300 in the carotid artery, a major vessel supplying blood to the brain, and continues to decrease deeper into the cerebral circulation[7]. Reynolds numbers smaller than 2300 indicate laminar flow, in which case the steady state assumption for CBF is well justified.

However, pulsatile pressure gradients can cause turbulent flow at lower Reynolds numbers. The Womersley number gives a ratio of inertial and viscous flow for unsteady or pulsatile flow.

Womersley numbers less than 12 indicate roughly laminar flow. The Womersley number is approximately four in the carotid artery[7] and less than one in smaller arteries and arterioles [8]. Bifurcations and stenosis can further lower the threshold for turbulence development. However, given the calculated Reynolds and Womersley numbers are relatively far from the turbulence threshold in the carotid artery, we assume that flow in the cerebral vasculature

#### Assumption 11.

*is laminar and can be approximated by Poiseuille flow*.

### C.4 Using the Lumped Parameter Assumption to Represent Cerebral Blood Flow Through a Vascular Tree

Poiseuille’s law (eq. 37) gives us an equation for average flow through a single straight tube. However, the cerebral vasculature is not just a single tube, but a complex vessel tree with structural morphology that varies between people and over time. The most comprehensive approach would be to map the entire cerebral vascular tree for each person, and calculate flow using either Navier-Stokes or Poiseuille’s law with boundary conditions. Unfortunately, this approach is nearly always intractable due to computational challenges and because mapping the entire human cerebral vasculature requires exceptional resources and is rarely, if ever, done.

In the pursuit of parsimony, if spatial components are not of primary interest to the specific application, or, more likely, unresolvable given technological or computational constraints, one approach is to assume that the entire vascular tree can be roughly approximated as a single tube. This assumption is justified by the following argument.

**Figure C.2:**
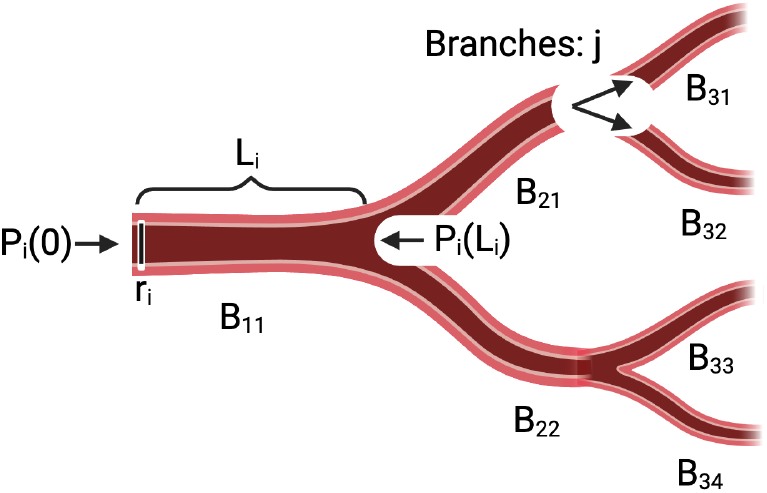
Toy example of branching vascular

Begin with a simplified vessel tree containing a single ‘parent branch’ (*B*_11_) directing blood into the skull that branches into ‘children’ branches (*B*_11*C*_ ∈ *{B*_21_, *B*_22_, *· · ·, B*_2*J*_ *}*). The children vessels subsequently branch into *N* layers (Fig. C.2). Each branch has radius and length (*r*_*ij*_ and *L*_*ij*_), where *i* refers to the layer and *j* refers to the branch in each layer.

First, define the ‘vascular’ 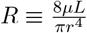 and rewrite eq. 37 as:

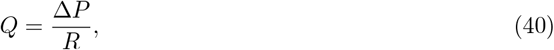

and assume

#### Assumption 12.

*pressure is continuous along the vessels and over bifurcations*.

Next, consider the first parent branch that bifurcates into *J* child branches. The pressure drop from the start of the parent branch to the end of the child branches is:

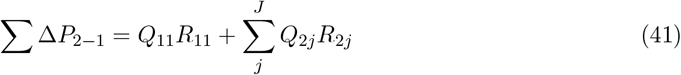

Assuming an incompressible fluid (assumption 1), steady state (assumption 6), the volumes of fluid in each branch are fixed, and fluid is not reflecting or ‘piling up’ on the bifurcations, the change in volume over time (e.g. the flow *Q*) must be conserved:

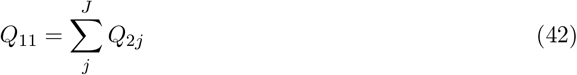

This assumption is not necessarily accurate for our system because radius changes in time. In subsequent sections, we argue changes in radius can be separated into slow and fast components, where the fast components is averaged over and the slow components are slow enough that this steady state assumption is generally valid.

Define the average flow across the system 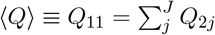 and rewrite eq. 41 as

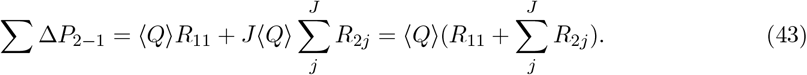

Define an effective resistance 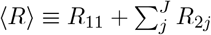. Now,

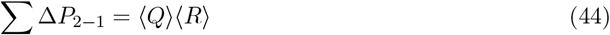

Recall Δ*P* is the gradient between pressure at the beginning of the vessel and end of the vessel. Assumption 12 allows the pressure at the beginning of a child vessel to be equal to the pressure at the end of its parent vessel. Further, using assumption 10, the outlet pressure is the same for all child vessels at the final layer *N* (in this case, *N* = 2).

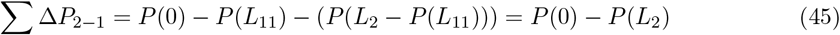

Therefore, ∑ Δ*P* is the gradient of the inflow and outflow pressures and the average flow across the vessel tree is:

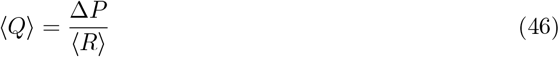

Recognizing that 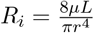,we can define ⟨ *R*⟩

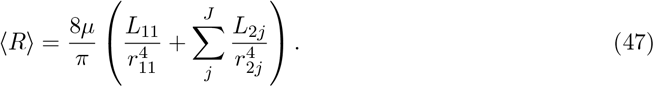

Now define an “effective” radius (⟨ *r*⟩) and length (⟨ *L*⟩) such that:

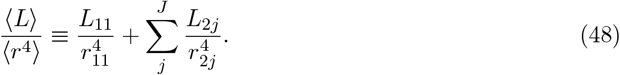

*The eventual goal of this model is to include physiological mechanisms that change effective radius. Since length and radius are coupled (eq. 48), in practice we fix length to isolate the effective radius*. The average flow across a parent and child vessel can be written in terms of effective geometry and a pressure gradient by substituting eq. 48 into eq. 46,

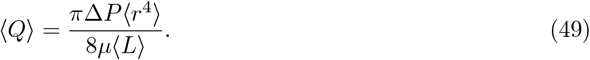

As seen here, pressure drop is proportional to the length, radius, and flow through the tube. Therefore, in larger vessel trees with heterogeneous geometry, the pressure drop through each branch will be heterogeneous and eq. 45 will not be for individual segments of the vascular tree. However, because the final arterioles at the end of the arteriole tree are assumed to have the same outlet pressure (ICP), eq. 49 can be used to model the full cerebral arteriole system by recursively repeating this process and redefining effective length and radius (eq. 48) with nested sums. Therefore, to model average flow through the complex vascular tree, we assume:

##### Assumption 13.

*Bulk dynamics the vascular tree can be roughly approximated as flow through a single tube*.

Some may recognize many of these arguments from Ohm’s and Kirchoff’s laws in electrical circuit theory. Because both are based on conservation laws, there are many parallels between fluid dynamics and electrical circuits, and it is common to argue fluid dynamics concepts using these parallels. This is called the hydraulic analogy. However, as seen above, the hydraulic analogy is not necessary to justify the bulk vessel assumption.

This argument neglects the impact of wave reflections and pulsatile pressure at vessel bifurcations. As discussed previously, there are multiple components ‘opposing’ flow or changes in flow, including the viscous and inertial terms in the Navier-Stokes equations and effects of boundary conditions. Instead of assuming (eq. 40), where resistance is quantified only by viscous forces 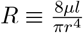,we could think of spatially averaged flow as simply:

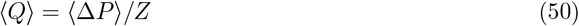

Here, *Z* is a scalar value that describes the bulk effect of the opposition to flow. This opposition to flow is called the “impedance”, because it describes how much the tube ‘impedes’ flow. Wave reflections contribute to the total impedance by absorbing energy. Rather than directly modeling these effects, we assume the vessel branching doesn’t change over the time scales we are modeling (hours) and adjust for vessel branching by estimating a general “impedance-type” term (see C.6).

### C.5 Accounting for Vessel Compliance By Modeling Average Flow Over Time

To obtain a model that can be simulated forward in time, we must take the derivative of eq. 49. We are interested in CBF dynamics over minutes to days. Pressure and radius change over these timescales. In terms of vessel geometry, we argue that:

#### Assumption 14.

*Changes in vessel length and viscosity are negligible compared to changes in radius on timescales of minutes to hours*.

This assumption is justified because vascular remodeling and angiogenesis typically occur on timescales of weeks to years, so *L* is roughly constant on shorter timescales. Changes in viscosity (*µ*) are also slow or dependent on medication administration and assume *µ* is constant (althrough interactions with medication could be a topic for future work). However, *r* changes over timescales of minutes to hours due to CVR. Further, *r* is raised to the power of four in eq. 21, therefore changes in *r* are dominant mathematically.

Now, let us rewrite eq. 49 explicitly stating temporal dependencies, grouping all constants into 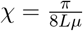,omitting spatial averaging notation (⟨·⟩) and rewriting Δ*P* as *P* for cleanliness:

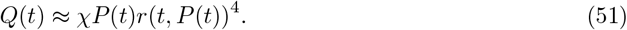

Taking the derivative of eq. 51 with respect to time (dropping dependencies for cleanliness), we obtain:

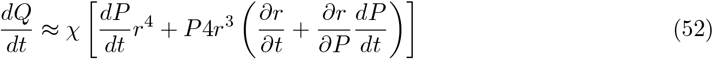

Conveniently, this expression distinguishes passive and active changes in the vessel radius, where passive changes are dependent on pressure 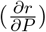 and active changes are dependent only on time 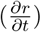.Notice that passive changes in 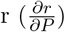 are scaled by changes in pressure 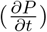.This supports the assumption:

#### Assumption 15.

*when pressure change is large and rapid*,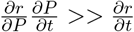

In the following section, we argue that passive and active changes usually occur on separable temporal scales and that averaging over short time scales allows us to isolate the effects of active changes in radius. To lay out an argument about when assumption 15 is relevant, we decompose pressure into its mean value 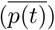 and fluctuations around that mean (*p*′(*t*)).

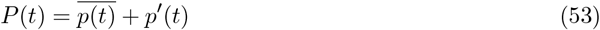

We can do a similar decomposition for radius to obtain:

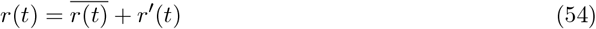

Since the Poiseuille equation is linear, we can also decompose flow using equations 53 and 55:

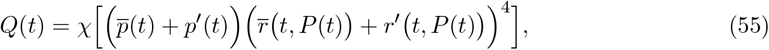

with corresponding derivative:

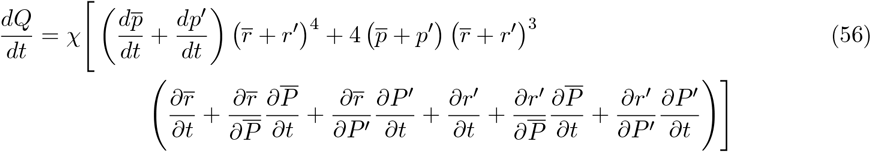

Most of the largest and fastest pressure changes are caused by rhythmic contraction and relaxation of the heart. These fast pressure changes typically oscillate around some mean value (which is often called mean CPP or mCPP). Therefore, we can assume that *p*′(*t*) describes pressure changes during the cardiac cycle and 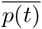 describes average blood pressure. Using assumption 15, we can further assume:

#### Assumption 16.

*over timescales shorter than multiple cardiac cycles (T <10 seconds)*,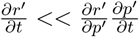

Alternatively, on longer time scales, the vasculature is very efficient at maintaining vessel radius via isometric contraction. As long as isometric contraction is functioning, we assume that:

#### Assumption 17.

*on longer timescales (longer than 10 seconds)* 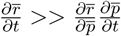.

Because CVR mechanisms are assumed to act on longer time scales [9], we assume that

#### Assumption 18.

*transient changes in pressure have no effect on active, slow change in radius* 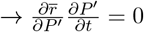.

Using these assumptions, we can rewrite eq. 56:

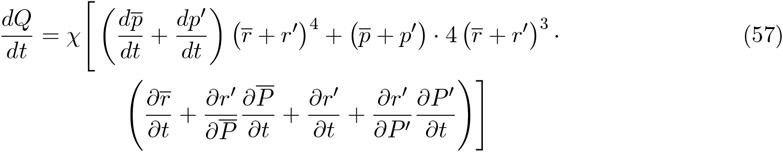

Blood flow measurements are typically limited to resolutions slower than one second. Moreover, average blood flow is generally more relevant in understanding if the brain is being adequately perfused than transient fluctuations in blood flow. Therefore, we will focus on the change in *average* blood flow over time. Define a smoothing operator:

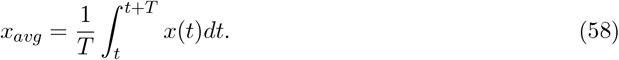

The change in average blood flow over time is 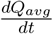.To calculate this, first calculate average pressure:

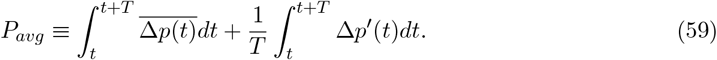

Because cardiac cycles (the heart beating) are roughly periodic, we make the assumption:

#### Assumption 19.

*Over short periods of time (T <* 10 *sec):* 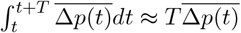 *and* 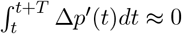.

Evaluating the eq. 60 given assumption 19 gives:

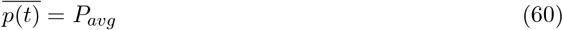

Similarly, changes in *r*′ are assumed to comprise passive changes due to rapid changes in pressure (assumption 15),

#### Assumption 20.

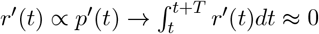

Smoothing over *Q* in equation 57 and including spatial averages for completeness allows us to write average blood flow as:

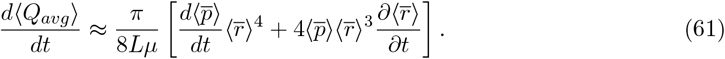

Here, 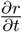 is not a constant and is controlled by mechanisms underlying CVR. Identifying a formula for 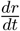 that allows us to estimate CVR function with patient data motivates the primary work of our study. Specifically, we construct and validate mathematical equations to control active vascular regulation based on physiological understanding of the myogenic, endothelial, and metabolic CVR mechanisms that contain functionality parameters that are estimable given our data. The overarching hypothesize of our paper is:

##### Hypothesis

Modeling active changes in vessel radius 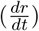 in the 0D lumped Poiseuille equation by mathematical representations of three CVR mechanisms will enable a model that can represent CBF dynamics.

### C.6 Estimating Scaling Factor *α*to Account For Vessel Bifurcations, Energy Loss, and Impaired Isometric Contraction

For our data, the magnitude of CBF is dependent on the location of the perfusion monitor. To obtain a scaling between personalized scaling between pressure and measured CBF for each patient, a scaling parameter *α* is estimated using the first 10 minutes of data. Because the value *L* in eq. 61 cannot be measured or defined external to radius, we replace *L* with *α* to act as a generalized scaling value between pressure and flow. This scaling very loosely captures multiple processes not previously accounted for: impedance and vessel compliance when isometric contraction is impaired.

Recall eq. 50 that argued for an estimated “impedance” that accounts for the total energy loss due to viscosity, bifurcations, etc. At a high level, this energy loss causes a smaller *Q* for a given *P*. We can account for this energy loss through the estimated scaling parameter *α* (although it is not exactly impedance because it does not fully account for the viscous forces when radius actively changes).

Estimated *α* also loosely accounts for effects of vessel compliance when isometric contraction is impaired. Assumption 17 allows us to prioritize active changes in vessel radius. This assumption is justified if isometric contraction is functioning because it reduces effects vessel compliance over long time scales. However, if isometric contraction becomes dysfunctional (assumed to occur when CVR is also dysfunctional and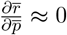), this assumption no longer holds. Therefore, in the case of impaired smooth muscle function,

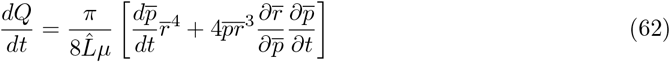

It can be seen that changes in pressure cause proportional changes in flow scaled both by *L* and by 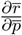.Therefore, as a crude approximation, we argue that estimating *L*, partially captures the effects of vessel compliance on flow in impaired isometric contraction. This approach has two key limitations. First, since *α* is not pressure dependent (unlike *r*), estimating *α* fails to capture temporal pressure dependent effects. Second, by approximating the effect of vessel compliance through *α*, we neglect it’s direct impact on *r*. Consequently, in the final digital twin, if isometric contraction is dysfunctional, while the estimated flow may still be reliable, the modeled radius is looses interpretability. To test this hypothesis, we assume isometric contraction may be impaired when CVR mechanisms are dysfunctional. In this case, *α* will be larger than when CVR mechanisms are functional. Despite these flaws, we feel that this simplification is pragmatically justified given our limited data set and prioritization of active vessel function over the arguably less interesting case where isometric contraction is impaired (less interesting because *Q* ∝ *P*).

### C.7 Further Justification for the Model From Windkessel Perspective

Instead of starting with Navier-Stokes or Poiseuille, many studies choose to model flow using what is known as the Windkessel model[10]. The Windkessel model analogizes blood flow to flow through the “Windkessel” in a fire-engine. Though a physical Windkessel is a chamber of compressible air, the analogy might be more readily visualized as a rigid tube attached to a balloon in such a way that fluid can flow through the tube or into the balloon to expand it. The total flow (*Q*) is the sum of flow through the tube (*q*_*R*_) or into the balloon (*q*_*C*_). Fluid will fluctuate between flowing into the balloon, through the tube, or both, depending on how much each component resists flow. Flow through the rigid tube is given by Poiseuille’s law, where total resistance to flow is written as 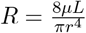,and

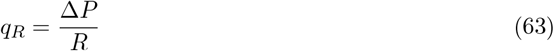

Alternatively, flow into the balloon is simply the increase of fluid volume in the balloon over time:

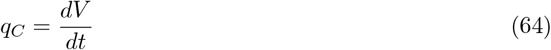

Recalling that we have measurements of the driving pressure, but not of the volume of flow into a hypothetical balloon, and that the volume is dependent on the driving pressure, we use the multivariate chain rule to express eq. 65 in terms of pressure:

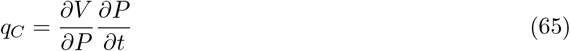

Using the same arguments from C.5, we can define balloon compliance as 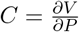.Therefore, the total flow through the system *Q* is:

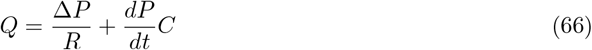

Not surprisingly (because they are all governed by the laws of physics) this Windkessel approach has many symmetries to Ohm’s Law for electrical circuits, where a resistor and capacitor represent characteristics of the vessel and can be aligned in series or parallel. Using this analogy, more complex spatial structure can be represented through multiple electrical circuits.

The Windkessel approach separates aspects of flow through a compliant tube into the ‘compliant’ component and the Poiseuille flow component. The primary question using this representation for our purposes is how to add active changes in vessel radius when both *R* and *C* have notions of vessel geometry that are separated mathematically but combined in the physical system.

However, one benefit of the Windkessel model is that the effect of compliance on flow is clearly seen to be dependent on the *rate of change* of pressure. When 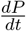 is very large, fluid is more likely flow into the balloon compared to when it is small. This concept is sometimes leveraged when trying to estimate *R* and *C* using pressure wave forms[11]. Specifically, it is assumed that during the cardiac contraction, in which 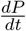 is very large, *q*_*c*_ *>> q*_*r*_, so 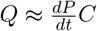,leaving just one variable to estimate (e.g. *C*).

In our situation, because 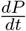 is the largest over short time scales *T* (as in section C.5), we can assume that the effects of vessel compliance will dominate on these short time scales. By averaging over these short timescales, we diminish the primary effects of compliance on the system. This dependence on 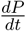 can be corroborated because it is known that the cerebral vessels display viscoelastic materials, even in the absence of smooth muscle (or implied dysfunction of smooth muscle)[12]. This argument helps justify “absorbing” vessel compliance into the estimated 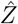 term.

Taking the integral of equation 66 and assuming

#### Assumption 21.

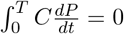,

we can define 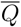 as the average flow over timescales longer than *T* to obtain:

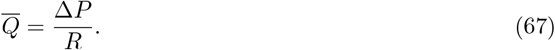

Assuming Poiseuille flow, we obtain:

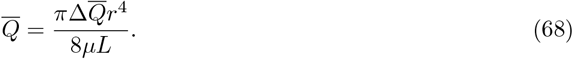

We can then make the same arguments as before to motivate the final equation 61 on which our model is based.

## Footnotes

1 also evidenced by strongly significant (P < 0.0001) univariate relationship between CMRO_2_ and PbTO_2_ in Rosenthal et al.[25]

2 patient CBF data was converted from ml/100g/min to ml/s (the units of simulated *Q*) by assuming average brain weight of 1200g[29]

3 This assumption may not hold in two cases for the cerebral vascular system. First, blood moves throughout the body. The cerebral vascular system is considered the portion of the vasculature that is located in or near the skull, but it is embedded and connected with the rest of the vascular system therefore, blood freely moves in and out of the cerebral vessels (e.g. to the heart) and mass may not be conserved. Second, in the case of hemorrhage, blood leaves the vessel but is still be located in skull.

4 This assumption includes the force of gravity

